# A century of theories of balancing selection

**DOI:** 10.1101/2025.02.12.637871

**Authors:** Filip Ruzicka, Martyna K. Zwoinska, Debora Goedert, Hanna Kokko, Xiang-Yi Li Richter, Iain R. Moodie, Sofie Nilén, Colin Olito, Erik I. Svensson, Peter Czuppon, Tim Connallon

## Abstract

Traits that affect organismal fitness are often very genetically variable. This genetic variation is vital for populations to adapt to their environments, but it is also surprising given that nature (after all) “selects” the best genotypes at the expense of those that fall short. Explaining the extensive genetic variation of fitness-related traits is thus a longstanding puzzle in evolutionary biology, with cascading implications for ecology, conservation, and human health. Balancing selection—an umbrella term for scenarios of natural selection that maintain genetic variation— is a century-old explanation to resolve this paradox that has gained recent momentum from genome-scale methods for detecting it. Yet evaluating whether balancing selection can, in fact, resolve the paradox is challenging, given the logistical constraints of distinguishing balancing selection from alternative hypotheses and the daunting collection of theoretical models that formally underpin this debate. Here, we track the development of balancing selection theory over the last century and provide an accessible review of this rich collection of models. We first outline the range of biological scenarios that can generate balancing selection. We then examine how fundamental features of genetic systems—including non-random mating between individuals, differences in ploidy, genetic drift, and different genetic architectures of traits— have been progressively incorporated into the theory. We end by linking these theoretical predictions to ongoing empirical efforts to understand the evolutionary processes that explain genetic variation.

## Introduction

Throughout nature—from birds, mammals and flies to flowering plants ^1–3^—individuals vary genetically in their ability to survive and reproduce, supplying the raw material for evolutionary adaptation. There are two broad schools of thought to explain this abundant genetic variation for fitness ^2,4^. One school proposes that selection continually removes genetic variation for fitness, while mutation continually replenishes it ^5^. Under this view, the resulting equilibrium between recurrent mutation and purifying selection (“mutation-selection balance”) accounts for fitness variation. The second school proposes that genetic variation for fitness is maintained by selection—a concept known as “balancing selection” ^6^. Balancing selection can arise from a variety of scenarios (Box 1), including selection favouring heterozygotes over homozygotes, selection for rare over common genotypes, and selective trade-offs in which genetic variants favoured in some contexts (*e.g.*, seasons, niches, sexes, life-history stages) are disfavoured in others.

### Box 1. Balancing selection and the concept of protected polymorphism

The concept of a “protected polymorphism” ^49^ defines the parameter conditions of a model that lead to balancing selection. The idea is simple: in single locus systems with two alleles segregating in an infinitely large population, polymorphism should be maintained if selection always favours the spread of each allele when it is rare. In the models outlined in Table 1, balancing selection occurs when the *A*_1_ allele (with frequency *p*) is favoured near frequencies *p* = 0, and the *A*_2_ allele (with frequency *q* = 1 – *p*) is favoured near frequencies *p* = 1. In other words, it occurs when the boundary equilibria (*p* = 0 and *p* = 1) are “unstable”. Conditions leading to this consistent rare-allele advantage (protected polymorphism, per Prout ^49^) are determined by stability analysis of the boundary equilibria ^192^.

**Table 1.** Single-locus models of balancing selection.

| | $A_1A_1$ | $A_1A_2$ | $A_2A_2$ |
| --- | --- | --- | --- |
| **Heterozygote advantage** | $1 - s_1$ | 1 | $1 - s_2$ |
| **Negative frequency-dependent selection** | $(1 - s_1 p)^2$ | $(1 - s_1 p)(1 - s_2)$ | $(1 - s_2)^2$ |
| **Meiotic drive** |  |  |  |
| Fitness of carrier | $1 - s$ | $1 - sh$ | 1 |
| $A_2$ transmission | 1 | $(1 + c)/2$ | 0 |
| **Fitness trade-off** |  |  |  |
| Fitness context 1 | 1 | $1 - s_1 h_1$ | $1 - s_1$ |
| Fitness context 2 | $1 - s_2$ | $1 - s_2 h_2$ | 1 |
| Average fitness | $1 - \frac{s_2}{2}$ | $1 - \frac{s_1 h_1 + s_2 h_2}{2}$ | $1 - \frac{s_1}{2}$ |

Table 1 outlines several scenarios that can lead to balancing selection (see Supplementary Material for stability analyses and conditions for balancing selection for each scenario). The parameters of these models have values between zero and one (*i.e*., 0 < *s*_1_, *s*_2_, *s*, *h*_1_, *h*_2_, *h*, *c* < 1), with *s*_1_, *s*_2_ and *s* representing homozygous fitness costs (*i.e.*, selection coefficients) of a particular allele and *h*_1_, *h*_2_ and *h* defining dominance coefficients. Selection coefficients are positive quantities that cannot exceed one (because fitness values cannot be negative), while dominance coefficients can range from partial-to-complete recessivity (0 ≤ ℎ < 0.5), to co-dominance (*i.e.*, ℎ = ℎ_1_ = ℎ_2_ = 0.5), to partial-to-complete dominance (0.5 < ℎ ≤ 1). The strength of meiotic drive (*c*, where 0 ≤ *c* ≤ 1) is scaled so that *c* = 0 corresponds to standard Mendelian segregation and *c* = 1 to 100% transmission of the drive allele.

In randomly mating diploid populations, heterozygote advantage generates balancing selection under all possible parameter conditions (*i.e.*, across the full “parameter space”; 0 < *s*_1_, *s*_2_ ≤ 1). However, the other scenarios in Table 1 generate balancing selection across a subset of the parameter space, with the remaining space favouring removal of one allele and fixation of the other (*i.e.*, directional selection). Assuming weak selection, allele frequency change over a single generation is approximately 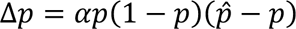 (eq. (1) in the main text), where 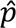 denotes the polymorphic equilibrium state where it exists. Balancing selection occurs when 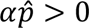 (the equilibrium *p* = 0 is unstable and *A*_1_ invades when rare) and 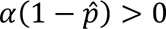 (the equilibrium *p* = 1 is unstable and *A*_2_ invades when rare), in which case the population evolves to the polymorphic equilibrium. For example, with heterozygote advantage and random mating, *α* = *s*_1_ + *s*_2_ and 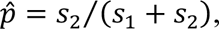 leading to 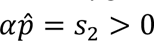 and 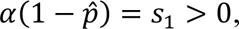 both of which must be true given that *s*_1_ and *s*_2_ are positive quantities. Expressions for *α* and 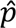 for the other scenarios are presented in the Supplementary Material.

Various scenarios of balancing selection cannot be accommodated by eq. (1). For example, in cases where balancing selection maintains more than two alleles at a locus, the frequency dynamics of each allele usually depend on all other alleles instead of just being proportional to *p*(1 − *p*). This is true of self-incompatibility systems, which can stably maintain many different alleles at a single locus, and also favour new alleles that may arise ^134,193–195^. Balancing selection may also lead to perpetual change rather than convergence to a polymorphic equilibrium, including cyclic dynamics in bi-allelic systems under strong negative frequency-dependent selection ^77^, and cycles in three-allele systems where the relative advantages of each allele relative to the others is nontransitive (*i.e.*, type A outperforms B, B outperforms C, and C outperforms A, as in rock-paper-scissors games; ^196^). Lastly, multiple polymorphic equilibria can occur in models with multiple loci, in single-locus models with sexually antagonistic selection ^68,69^, or with non-linear frequency-dependent selection, such as game theory models involving interactions between more than two individuals ^197,198^.

Despite considerable debate during the second half of the 20^th^ century ^4^ and recent renewed interest in balancing selection (Fig. 1A), the contribution of balancing selection to the maintenance of fitness variation remains unresolved ^2^. Little more than a decade ago, however, the prevailing view was that balancing selection was probably of minor importance in evolution. Summarising the sentiment, Asthana et al. ^7^ wrote that “balancing selection […] has not been a significant force in human evolution”, while Hedrick ^8^ similarly suggested that “a low proportion of loci in the human genome are under long-term balancing selection”.

**Figure 1.**
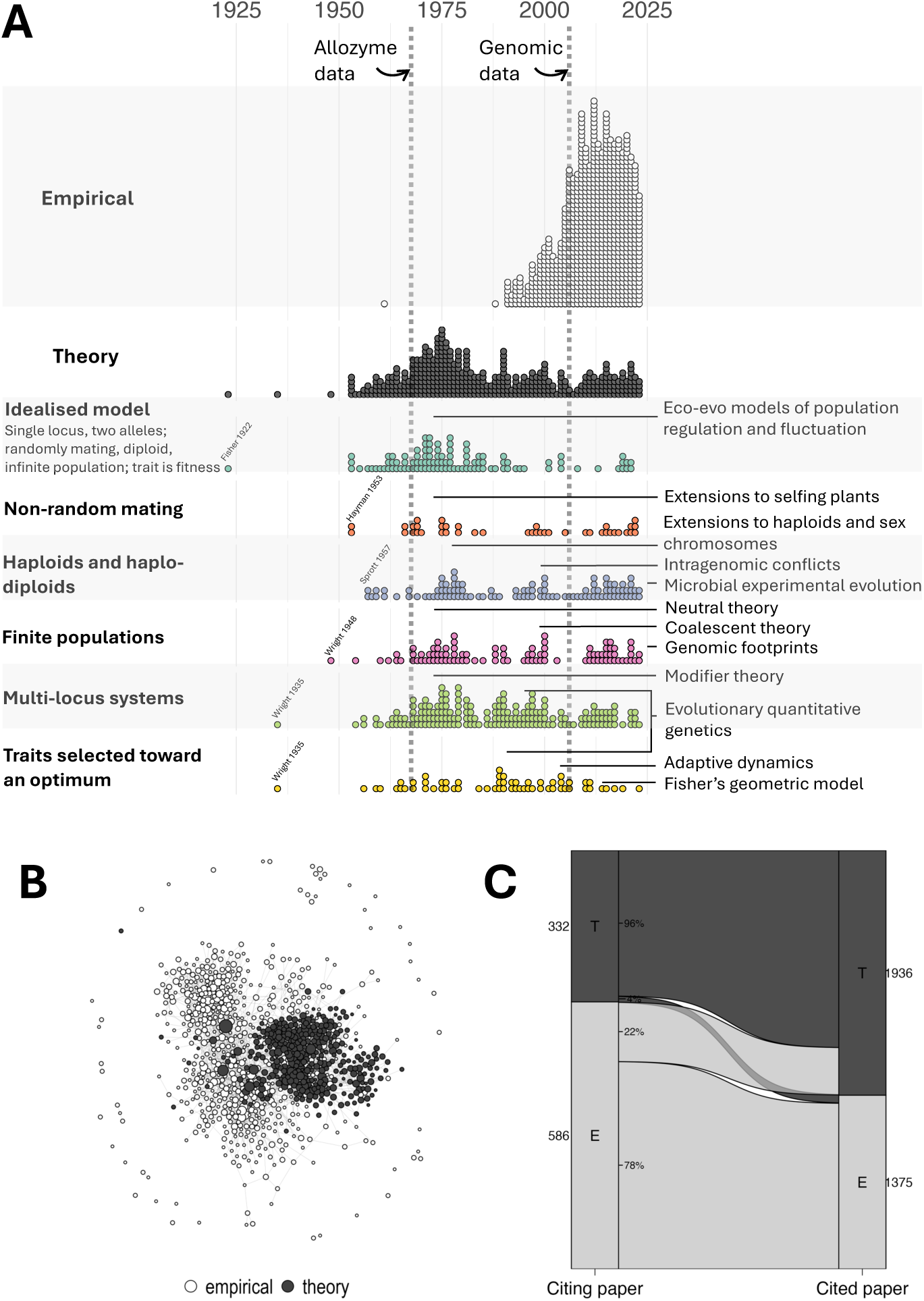
**A.** A timeline of balancing selection theory and data. Each dot represents a paper on balancing selection. Papers were identified by a targeted search of the Web of Science, followed by manual curation of theory papers missed by the search (N = 872 empirical papers, N = 402 theory papers; see Supplementary Material for details of the collection methods). The top panel illustrates empirical research on balancing selection, which grew dramatically in the 1980s, spurred by the discovery of sequence variants (protein polymorphisms termed allozymes) and DNA variants, including molecular markers (e.g., microsatellites) and, later, genome sequences. The next panel illustrates the gradual growth of balancing selection theory, which spiked in the mid-1970s, spurred by the debate over neutral theory and a desire to account for abundant protein polymorphisms in natural populations ^4,15,22,48^, and has gained ground in the genome era (2010s). The bottom panels zoom in on the theoretical papers and separate them into different aspects of biological complexity (left), following the structure outlined in ‘Adding biological complexity to models of balancing selection’. Individual theory papers can span multiple aspects of complexity at once. The lines (right) highlight some particularly influential developments in the theory. **B.** Citation network of theory and empirical papers. Each node (circle) represents one article in the dataset. Node colours denote article types (empirical, theory); node size is proportional to number of times the article is cited locally within the network (*i.e.*, not the global citation count); edges show citations within the local network. Unconnected nodes do not cite articles within the network and are not cited by articles within the network. **C.** Citation patterns for each article type (E = empirical, T = theory) within the local network. Box and flow widths reflect the number of articles of a given type.

Why did this become the prevailing view? First, few empirical demonstrations of balancing selection were clearly documented at the time. The most prominent examples— including beta-globin alleles in humans ^9^, self-incompatibility alleles in outcrossing plants ^10,11^, chromosomal inversion polymorphisms in *Drosophila* ^12^, and the major histocompatibility complex alleles in vertebrates ^13^—were decades-old and viewed as outliers. When early genome scans failed to detect polymorphisms under long-term balancing selection ^7,14^, the view that balancing selection was exceedingly rare was reinforced. Second, balancing selection hypotheses faced several theoretical objections. In particular, ubiquitous balancing selection on the abundant polymorphisms discovered in natural populations ^15,16^ seemed unlikely given the exorbitant mortality cost (“genetic load”) implied by such a hypothesis ^15,17^, though ecological counterarguments soon tempered this criticism ^18–23^. Meanwhile, the intuition that genetic trade-offs often generate balancing selection was undermined by theory demonstrating that trade-offs typically result in loss, rather than maintenance, of genetic polymorphism ^24,25^. Finally, models of quantitative traits selected towards an intermediate optimum (*i.e.*, traits under “stabilizing selection”) highlighted restrictive conditions for persistent balancing selection at the genetic loci underlying the trait ^26–28^. The notion that traits are often quantitative and selection is often stabilizing ^29^ seemed, therefore, to minimise the role of balancing selection.

Recent advances cast doubt on this consensus. New methods for detecting balancing selection in genomic sequences have now uncovered hundreds of candidate balanced polymorphisms ^30,31^, in contrast to the small number of pre-genomic ones. Although most candidates require further validation, the current list likely represents the tip of the iceberg, given that genomic methods for inferring balancing selection are often under-powered and rely on long-term signals ^31^. Second, arguments about genetic loads have become less relevant to contemporary debates about balancing selection because genome-wide genetic diversity is now thought to be predominantly neutral or mildly deleterious ^32,33^. The focus has instead shifted to the role of balancing selection in maintaining the extensive genetic variation for life-history traits and other fitness components, which is indeed too high to be explained by mutation alone ^2,3,34,35^. Finally, new developments in theoretical modelling of adaptation give prominence to balancing selection. While earlier models emphasised the often-restrictive conditions for maintaining polymorphism indefinitely, newer models have highlighted the potential for short-lived episodes of balancing selection during the evolution of traits toward their optimum ^36–38^. Moreover, a surge in theoretical models of fluctuating selection—inspired by recent evidence of predictable seasonal fluctuations of nucleotide polymorphism in *Drosophila* ^39–41^ and other species ^42^—have highlighted broader conditions for balancing selection than implied by classical models ^42–45^.

The renewed interest in balancing selection has prompted several reviews of empirical progress in detecting it ^13,30,31^, but a review of the underlying population genetic *theories* of balancing selection is lacking. Here, we present a comprehensive and accessible overview of this theory. As is true for most models, the devil is in the details, and a deeper understanding requires some engagement with the particulars. We therefore provide a largely verbal overview in the main text, collect technical aspects in Boxes, and elaborate further in mathematical derivations presented in the Supplementary Material.

Balancing selection has a long history of study over the last century (Fig. 1A). We begin our review with the idealised conditions that characterise the very first models of balancing selection: *i.e.*, single biallelic genetic loci evolving in large, randomly-mating, diploid populations. These conditions mirror Fisher’s ^46^ classic analysis of heterozygote advantage ^47^. As the 20^th^ century progressed, the major technical innovations and debates that stimulated empirical evolutionary biology (*e.g.*, the advent of allozyme and genomic data; Fig. 1A) coincided with the development of theories of balancing selection. We therefore examine how various aspects of biological complexity were gradually incorporated into the theory (Fig. 1A), including non-random mating, deviations from diploidy, genetic drift, linkage and recombination between loci, and different forms of trait inheritance and phenotypic selection. We close by outlining progress in linking the theoretical predictions about balancing selection to empirical data (Fig. 1B,C).

### Models of balancing selection: the basics

#### The concept of balancing selection

Balancing selection occurs when natural selection—without the intervention of other processes such as genetic drift, mutation, or migration—maintains genetic polymorphism. It can arise from a variety of scenarios (outlined in Box 1), but in all scenarios rare alleles have an advantage over common alleles. This aligns with the intuition that balancing selection maintains variation by opposing the loss of rare variants, as well as its operational definition of “protected polymorphism” in mathematical population genetics (^49^; see Box 1). Still, this definition is easy to misinterpret, and we outline some possible misconceptions in Box 2.

##### Box 2. Clarifying misconceptions about balancing selection

The definition of balancing selection is well established in theoretical population genetics (Box 1) but is easy to misinterpret. We try to clarify some possible misconceptions below.

- ***It is not a category of trait selection.*** Natural selection refers to the differential survival or reproduction of individuals expressing different phenotypes, which can occur regardless of the genetic basis of the trait in question. In contrast, balancing selection— a concept from population genetics—is defined by the *genetic consequences* of natural selection. Specifically—in the absence of other evolutionary factors—it refers to any form of phenotypic selection that maintains polymorphism at one or more loci within the genome (Box 1). Thus, different forms of trait selection (*e.g.*, directional, disruptive, or stabilizing; ^29^) potentially generate balancing selection at trait-affecting loci ^27,154^, but balancing selection does not necessarily accompany a given form of trait selection.
- ***It is not equivalent to heterozygote advantage, negative frequency-dependent selection on genotypes, or trade-offs.*** Each of these scenarios can generate balancing selection, but they need not. For example, heterozygote advantage does not always lead to balancing selection in haplodiploids and inbreeding populations ^92,107^ (Fig. 3), nor do certain models of negative frequency-dependent selection ^199^, nor do many trade-offs (Box 3). However, balancing selection *does* imply that the “marginal fitness” of each *allele* declines with its frequency (Fig. 2B). This type of frequency-dependence should not be confused with the more common usage of the term (*i.e.*, that the fitness of each *genotype* declines with its frequency)—indeed it applies, for example, to heterozygote advantage (Fig. 2B), in which the fitness of each genotype is independent of frequency.
- ***It is not necessarily a long-term process.*** Balancing selection is often used interchangeably with long-term balancing selection, possibly because genome scans often rely on long-term rather than short-term signals (*i.e.*, long-term signals have durations greater than 4*N_e_* generations, where *N_e_* is effective population size; ^31^) or because the term “stability” (Box 1) is taken to imply perpetual stability. However, timescale is not part of the definition of balancing selection. It can be transient (*e.g.*, over relatively few generations) or persistent (*e.g.*, over many thousands of generations), with each scenario leaving a different genomic footprint.
- ***It does not always elevate polymorphism in real populations.*** When drift is strong relative to selection, balancing selection can lead to patterns of polymorphism that are indistinguishable from neutrality, both at the target of selection or at linked loci. It can even *reduce* levels of genetic diversity, relative to neutrality, when the equilibrium frequency of the selected allele is close to zero (see section: ‘Finite populations’).
- ***It is not the only evolutionary process that “maintains” genetic polymorphism.*** Balancing selection maintains polymorphism in the absence of other evolutionary factors. However, interactions between mutation, migration, genetic drift, and positive or purifying selection are also capable of maintaining polymorphism ^52,200,201^. For example, purifying selection against harmful genetic variants is offset by mutation, which continuously introduces new harmful variants. Genetic variation can then be maintained at the equilibrium between these two opposing processes (“mutation-selection balance”).

#### Scenarios of balancing selection

Consider an idealised model in which a single genetic locus segregates for two alleles, the population is infinitely large and diploid, mating is random, and generations are discrete. One scenario that generates balancing selection under these assumptions is heterozygote advantage (*i.e.*, overdominance), in which heterozygotes have higher fitness (*i.e.*, higher survival and/or fertility) than homozygotes (Fig. 2A) ^46,50^. Each allele, when rare, is expected to increase in frequency because rare variants are predominantly found in (fit) heterozygotes, whereas common variants are mostly found in (less fit) homozygotes. Although the fitness of each genotype is independent of its frequency, the average fitness of each *allele*—its so-called “marginal fitness”—declines with the allele’s frequency (Fig. 2B). Under heterozygote advantage, the population evolves towards a polymorphic equilibrium at which the marginal fitness of both alleles is equal (Fig. 2C). Heterozygote advantage can also maintain more than two alleles at a single locus, though conditions for maintenance of many alleles become restrictive unless there is a high degree of symmetry among homozygous and heterozygous genotypes for the set of alleles ^51,52^.

**Figure 2.**
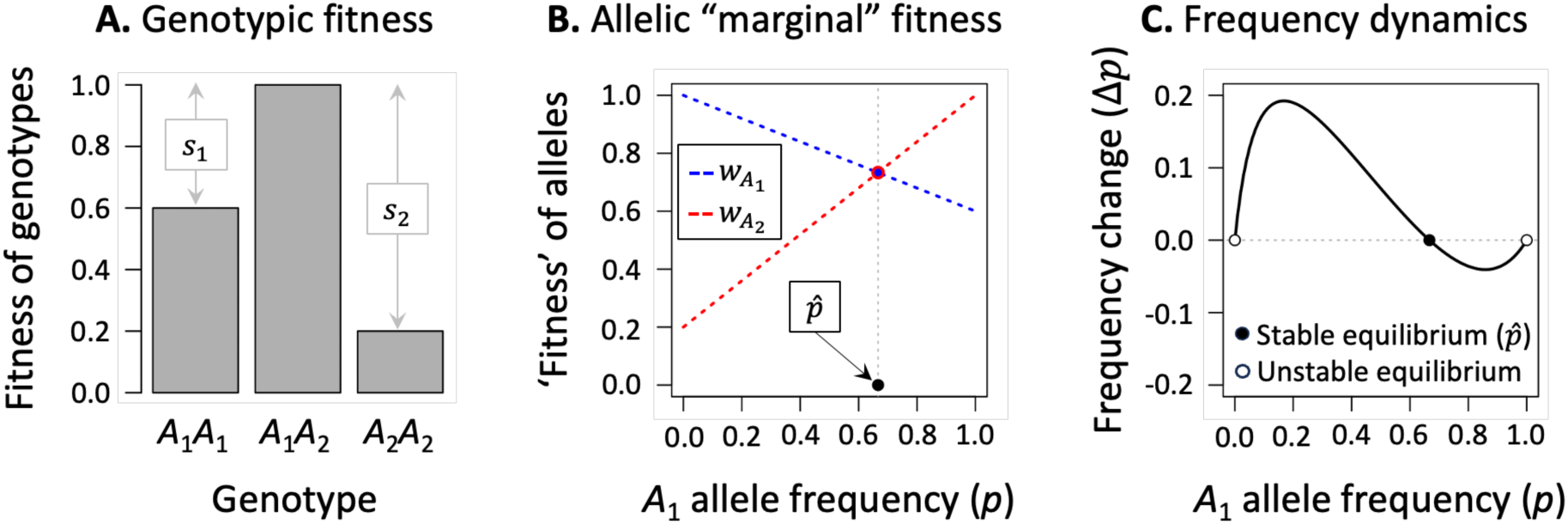
Heterozygote advantage as an example of balancing selection. **A.** For a locus with two alleles (*A*_1_ and *A*_2_, at frequencies *p* and *q*, respectively; Box 1), the fitness of *A*_1_*A*_1_ homozygotes declines by *s*_1_, and *A*_2_*A*_2_ declines by *s*_2_, relative to heterozygotes (fitness of the best genotype is scaled to one). **B.** Though fitness per genotype is frequency-*in*dependent, the average transmission rate of each allele to the next generation (its ‘marginal fitness’) is frequency-dependent. In outbred, randomly mating populations, the marginal fitness of the *A*_1_ and *A*_2_ alleles (*w*_*A*1_ = 1 − *ps*_1_and *w*_*A*2_ = 1 − (1 − *p*)*s*_2_, respectively) are each negative frequency-dependent (the marginal fitness of each allele declines with the frequency of that allele), becoming equal at the polymorphic equilibrium 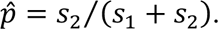 **C.** Whether the *A*_1_ allele increases (Δ*p* > 0) or decreases in frequency (Δ*p* < 0) depends on its current frequency relative to the equilibrium.

Balancing selection is also likely, though not inevitable, when the fitness of each *genotype* declines with its frequency in the population—a scenario referred to as negative frequency-dependent selection ^53,54^. In such cases, rare genotypes may consistently experience fitness advantages over more common ones. There is a vast ecological literature on negative frequency-dependent selection ^55^ and myriad ways to model it (see chapter 5 of ^56^; ^57,58^). In Box 1, we present a simple illustrative example that can either lead to balancing selection, or to fixation of one allele and extinction of the other (*i.e.*, to “directional selection”; see Supplementary Material).

Balancing selection can also arise from a variety of genetic trade-offs, in which alleles that are advantageous in some contexts are harmful in others (Box 1). In the case of meiotic drive, a “driver” allele has a transmission advantage because it finds itself in more than half of the gametes produced by heterozygotes. However, driver alleles may also lower the fitness of individuals that carry them ^59,60^. The trade-off between an allele’s transmission advantage and its fitness cost to carriers can then potentially generate balancing selection (Box 3). Trade-offs can also arise between temporally fluctuating environmental conditions ^42–45,61–63^, different resources or “niches” used by the population (“niche antagonism”; ^64–67^), females and males (“sexually antagonistic selection”; ^68–70^), and life-history traits (“antagonistic pleiotropy” between *e.g.,* survival and fertility; ^71,72^).

##### Box 3. Dominance and opportunities for balancing selection through trade-offs

Trade-offs can generate balancing selection, but they often will not ^25^. The following figure outlines conditions for balancing selection under trade-off scenarios where a polymorphic equilibrium is attainable, *i.e.*: meiotic drive (MD), antagonistic pleiotropy (AP), sexually antagonistic selection (SA), and niche antagonism (NA).

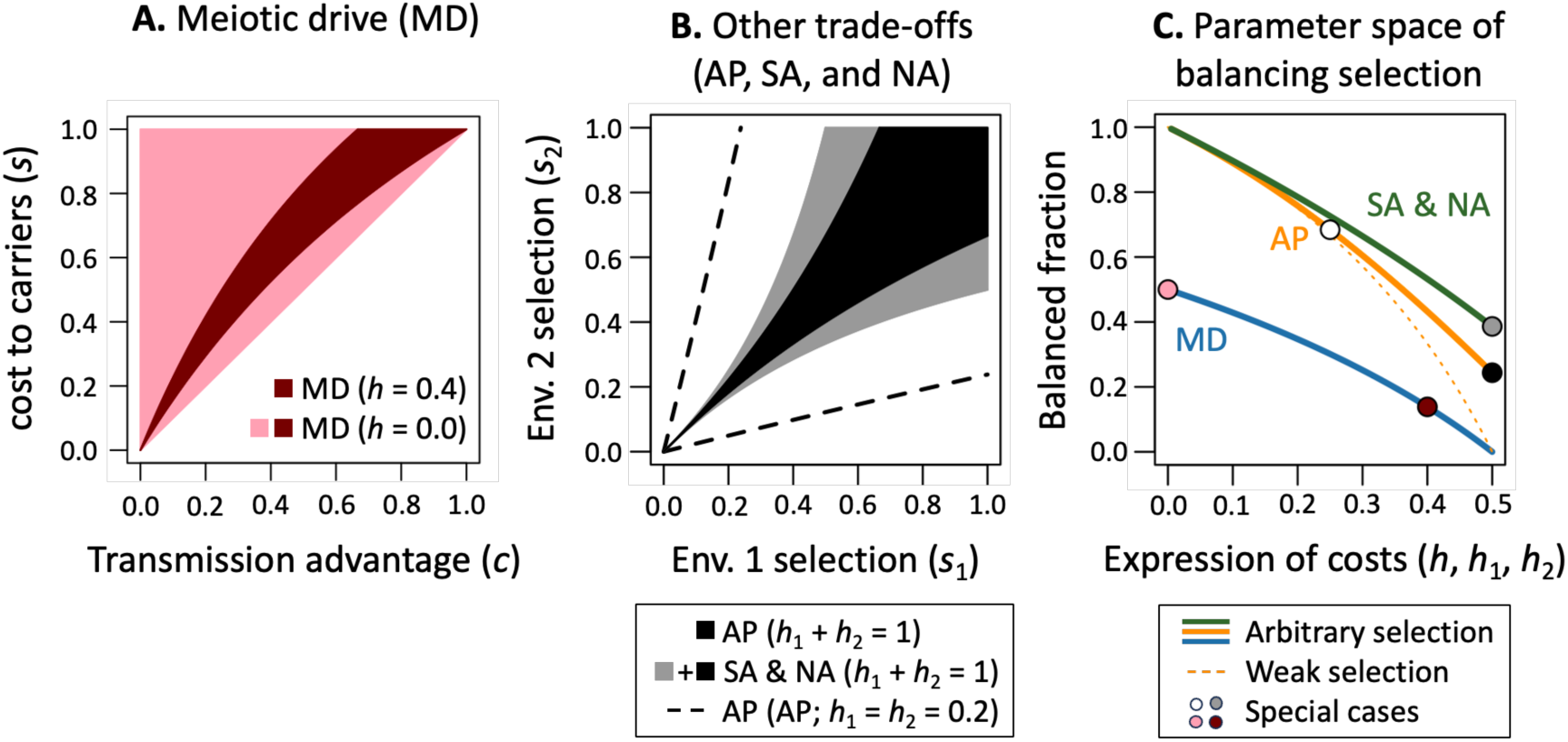

Balancing selection is possible under MD (shaded regions of panel A; ^60^) when the homozygous fitness cost (selection coefficient, *s*) of a drive allele is greater than its segregation advantage (*c*), and the cost is at least partially masked in heterozygous carriers (*h* < 0.5; parameters as defined in Box 1). Other trade-off scenarios can generate balancing selection when fitness costs are co-dominant on average (*h_1_, h_2_* = 0.5, shaded regions of panel B; ^24,60,74^), though conditions are restrictive unless selection is strong. Conditions for balancing selection become more permissive when the expression of each allele is recessive in contexts where it is costly and dominant in contexts where it is beneficial ^48,52,73–76^ (*h_1_, h_2_* < 0.5). In the case of AP, SA and NA, this is known as a “dominance reversal” (or, more specifically, a “favourable reversal of dominance”).

Dominance reversals increase the likelihood that the average fitness of the heterozygotes is greater than the average for homozygotes (*i.e.*, that “net heterozygote advantage” arises across contexts of selection). In Table 1, within Box 1, a net heterozygote advantage occurs when *s*_1_ℎ_1_ + *s*_2_ℎ_2_ < min(*s*_1_, *s*_2_). In the extreme case of a complete dominance reversal (*h*_1_ = *h*_2_ = 0), a net heterozygote advantage and balancing selection will arise across the entire parameter space for *s*_1_ and *s*_2_. A net heterozygote advantage is a requirement for balancing selection through AP ^72^, but not under SA or NA (*e.g.*, co-dominant costs do not lead to net heterozygote advantage but permit balancing selection; ^69,74^).

The NA model considered above consists of two equally sized ecological niches with random dispersal and “soft selection”, which is a special case of Levene’s more general model ^64,65^. Balancing selection through NA becomes less likely when one niche is more common than the other and selection is “hard” (^62^; see Supplementary Material). Conditions become more permissive when gene flow is limited between patches (*e.g.*, due to habitat selection, assortative mating by niche type, and extrinsic barriers to gene flow; ^65^). Broader mathematical criteria for balancing selection are described in the Supplementary Material.

In each trade-off model, the balance between benefits and costs must be just right to maintain the polymorphism (Box 3), and conditions leading to directional selection are often more permissive than those leading to balancing selection ^24,25^. Conditions for balancing selection are particularly restrictive when selection is weak and the benefits and costs associated with each allele have similar degrees of dominance (Box 3). However, conditions for balancing selection become substantially more permissive when the cost of expressing each allele is at least partially recessive while its benefit is dominant (Box 3). Such “dominance reversals” ^48,52,73–76^) can lead to a “net heterozygote advantage”, in which the mean fitness of heterozygotes is higher than the mean fitness of homozygotes. Net heterozygote advantage is a sufficient condition for balancing selection in randomly mating diploid populations (see Box 3 for elaboration).

#### Allele frequency dynamics under balancing selection

Although the details of individual balancing selection scenarios vary, their allele frequency dynamics often simplify to a common form that highlights their conceptual unity (Box 1). For models with a single polymorphic equilibrium, allele frequency change per generation is approximately:

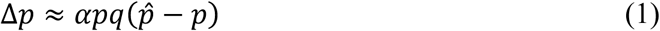

where *p* and *q* = 1 – *p* refer to *A*_1_ and *A*_2_ allele frequencies, *α* is the net strength of selection (here assumed to be weak, such that |*α*| ≪ 1), and 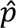 is the polymorphic equilibrium. Under balancing selection, *α* > 0 and 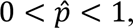 with values of *α* and 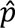 depending on the scenario of balancing selection and its underlying parameters (Box 1 and Supplementary Material).

Several insights emerge from eq. (1). First and most obviously, evolutionary change requires genetic variation at the locus (0 < *pq* < 1). Second, given that *αpq* must be positive when genetic variation is present, the term 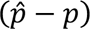 determines the direction of evolutionary change. The frequency of *A*_1_ increases when it is below the equilibrium, decreases when it is above the equilibrium, and remains stable at the equilibrium. Therefore, in the absence of genetic drift, balancing selection causes a genetically variable population to evolve towards the polymorphic equilibrium. This is illustrated in Fig. 2C for the case of heterozygote advantage, though similar dynamics characterise each of the scenarios outlined in Box 1.

While most models of balancing selection can be recast in the form described by eq. (1), there are notable exceptions. For example, strong negative frequency-dependent selection^77^ and/or temporal fluctuations in the direction of selection can maintain polymorphism and exhibit predictable patterns ^43,63,78^, but there is no polymorphic equilibrium state that the population eventually reaches. Models with multiple alleles ^51,79^, or multiple polymorphic equilibria ^68,69^, also cannot be described by eq. (1) (Box 1).

### Adding biological complexity to models of balancing selection

All models are simplifications, and the scenarios of balancing selection presented above obviously omit important aspects of biological complexity. In some cases, violations of their simplifying assumptions can fundamentally alter opportunities for balancing selection, or the evolutionary stability of balanced polymorphisms. For example, while many populations are indeed diploid and mate nearly randomly at most loci, others—such as haplo-diploid species, or self-fertilising plants—are not. Conditions for balancing selection that go beyond the diploid and randomly-mating ideal were increasingly explored during the second half of the 20^th^ century ^80,81^ (Fig. 1A). Likewise, while models that ignore genetic drift are often useful approximations of reality, drift is an inevitable feature of real populations with the potential to destabilise balanced polymorphisms. The rise of the neutral theory of molecular evolution ^82^, coalescent theory ^83^, along with the increasing feasibility of computer simulations, led to a burst of finite-population models of balancing selection (*e.g.*, ^84–86^). Finally, while single-locus models may well describe traits with a simple genetic basis (*i.e.*, whose expression is dictated by few genetic variants), they apply awkwardly to continuous traits. The advent of “modifier” models, which focus on multi-locus genetic systems (*e.g.*, ^87,88^), and various models that link genotype, phenotype and fitness (*e.g.*, ^27,36,89–91^) extended the scope of theories of balancing selection beyond single loci (Fig. 1A). We outline the consequences of each of these aspects of biological complexity below.

#### Non-random mating

Mating patterns influence the proportion of heterozygotes, and thereby opportunities for balancing selection in scenarios where heterozygotes are favoured over homozygotes (*i.e.*, in cases involving true or net heterozygote advantage; see Boxes 1 and 3). Mating patterns that decrease the proportion of heterozygotes (*e.g.*, inbreeding or positive assortative mating by genotype) tend to decrease the range of conditions leading to balancing selection ^80,92^, while mating patterns that increase the proportion of heterozygotes (*e.g.*, disassortative mating by genotype) tend to do the opposite ^93,94^.

Consider the widespread case of inbreeding, which reduces heterozygote proportions relative to random mating. Models of balancing selection with inbreeding can be expressed using eq. (1), but with parameters *α* and *p*^ adjusted to include the population’s inbreeding coefficient (*F*) where *F* = 1 denotes complete inbreeding and *F* = 0 denotes complete outcrossing. Under weak heterozygote advantage, the net strength of selection becomes *α* = (1 − *F*)(*s*_1_ + *s*_2_), the polymorphic equilibrium is 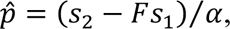 and balancing selection arises when *Fs*_1_ < *s*_2_ < *s*_1_⁄*F*. Thus, inbreeding decreases the parameter range for balancing selection by a factor of 1 – *F* when selection is weak (Fig. 3A), and somewhat less when selection is strong ^95^. Inbreeding also depresses the scope of balancing selection resulting from trade-offs ^80,96–99^. This can arise when there is net heterozygote advantage (*e.g.*, sexually antagonistic selection with dominance reversal ^100^), or when self-fertilization reduces the strength of selection through the male sex function of hermaphrodites, which promotes the fixation of sexually antagonistic alleles that benefit females ^92,98,101^. By contrast, inbreeding has little effect on balancing selection arising from negative frequency-dependent selection ^92,102^.

**Figure 3.**
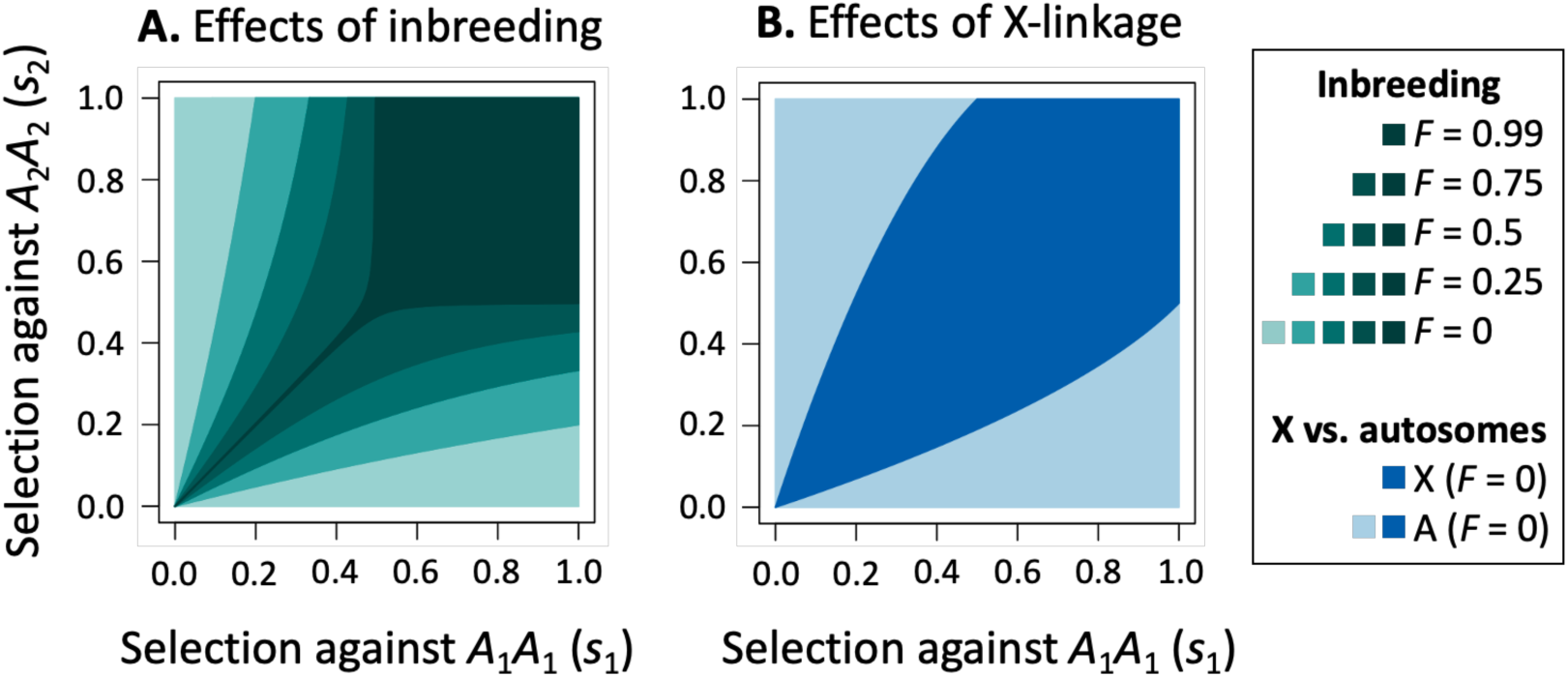
Consequences of inbreeding and X-linked (or haplo-diploid) inheritance on the maintenance of polymorphisms under heterozygote advantage. Whereas outbred and diploid populations can maintain a balanced polymorphism over the entire parameter space of selection (the full range of values for *s*_1_ and *s*_2_; 0 < *s*_1_, *s*_2_ ≤ 1), inbred diploid populations (**A**) and X-linked genes or haplo-diploid populations (**B**) reduce the parameter space for balanced polymorphism (dark shaded regions). Results are based on exact criteria for balancing selection ^107,108^.

Even when balancing selection is predicted to occur, inbreeding reduces the effective size of the population ^103^ and increases selective interference between genetically linked loci ^104^. This promotes the loss of polymorphism by reducing the efficacy of balancing selection relative to genetic drift ^92^ (see section: ‘Finite populations’). On the other hand, inbreeding can sometimes facilitate balancing selection of multi-locus allele combinations ^101^ and intensify population genomic signals of long-term balancing selection ^105,106^.

#### Haploids and haplo-diploids

Even if heterozygotes have higher fitness than homozygotes, such advantages become irrelevant in predominantly haploid populations, haploid individuals, or haploid stages of a life cycle (*i.e.*, gametes or the gametophyte stage of plants). In fully haploid populations, there is no scope for maintaining polymorphism by heterozygote advantage, meiotic drive, or antagonistic pleiotropy. Balancing selection may still arise from negative frequency-dependent selection ^109,110^ or from trade-offs between niches, seasons or sexes ^111–114^. However, the conditions for balancing selection under such trade-offs are highly restrictive. In the case of trade-offs between sexes or niches, balancing selection conditions under haploidy are equivalent to those under diploidy with co-dominance (Box 3).

When populations experience selection in both diploid and haploid life stages, or when diploid and haploid individuals co-exist, conditions for balancing selection tend to be intermediate between purely diploid or haploid populations ^107,115^. For example, at loci where one sex is haploid and the other is diploid (*e.g.*, haplodiploids, X-linked genes that are hemizygous in males), heterozygote advantage in the diploid context favours the maintenance of polymorphism, while the haploid context typically favours its loss. Hence, balancing selection only occurs when heterozygote advantage in the diploids outweighs directional selection in the haploids (as in the dark shaded region of Fig. 3B, where *s*_1_ and *s*_2_ are similar in magnitude, implying weak directional selection in haploids). In the rest of the parameter space, the allele that is least harmful in homozygous or haploid individuals is fixed (light shaded regions of Fig. 3B). Conditions for X-linked balancing selection are also reduced in models of female meiotic drive, antagonistic pleiotropy, and trade-offs between niches (Supplementary Material; ^116^).

Although X-linked inheritance usually restricts conditions for balancing selection, exceptions can occur. Cases of male meiotic drive are especially complex because they are associated with skewed sex ratios, altered population size dynamics, and potentially extinction ^117–119^. In models of sexually antagonistic selection ^73,120,121^, the sex-specific dominance coefficients of the alleles determine whether polymorphisms are more readily maintained at autosomal (diploid) or X-linked loci. Specifically, the X is more permissive for balancing selection when male fitness costs of sexually antagonistic alleles show moderate to strong dominance (*h_1_* > 1/(2 – *s_1_*), where *h_1_* and *s_1_* represent male dominance and selection coefficients). Autosomes are more permissive otherwise ^122^. Comparable results arise under “ploidally antagonistic selection”, where different alleles are favoured in the haploid and diploid stages of a life cycle ^115^.

#### Finite populations

Real populations are finite and subject to genetic drift, which causes random deviations from deterministic evolutionary trajectories. Despite this randomness, population genetic models generate clear predictions for the stationary (*i.e.*, long-run) probability of observing each possible allele frequency state, as a function of the effective population size (*N_e_*) and the specific scenario of balancing selection (Fig. 4A). For the scenarios in Box 1, the stationary distribution for the *A*_1_ allele is:

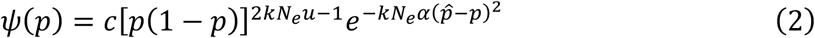

**Figure 4.**
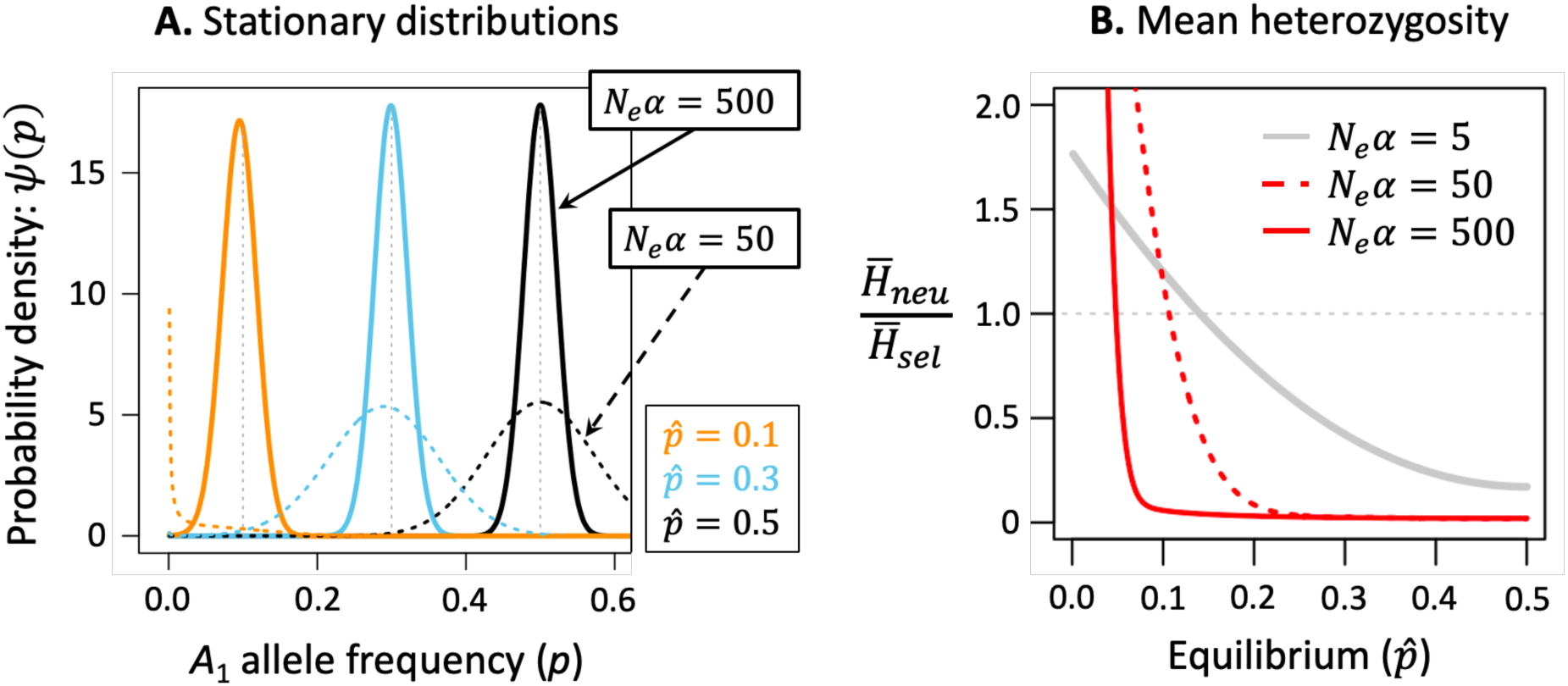
Interactions between balancing selection and genetic drift. **A.** Diffusion approximations are used to predict stationary (long-term) distributions (eq. (2)) for three equilibrium frequency states, and two strengths of selection. **B.** The ratio of mean genetic (*e.g.*, nucleotide) diversity for neutral loci relative to loci under balancing selection is shown. For each parameter combination, the expected heterozygosity is numerically calculated as 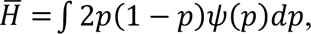 where 2*p*(1 – *p*) is the heterozygosity and 𝜓(*p*) is the stationary distribution. When the ratio is greater than one, then expected heterozygosity is higher under neutrality than under balancing selection. Results for both panels assume 2*kN_e_u* = 0.01 and *k* = 2 (diploidy).

where *α* is the net strength of selection (assumed to be weak), *k* is the number of gene copies carried by each member of the population (*k* = 2 for diploids and *k* = 1 for haploids), *u* is the mutation rate per locus (assumed to be the same for each allele), and *c* is a constant that ensures that the distribution integrates to one.

Analyses of the stationary distribution reveal two important consequences of drift for polymorphisms under balancing selection. First, given no additional mutations entering the population, a balanced polymorphism will eventually be lost despite selection to maintain it (see p. 165 of ^123^). Second, and somewhat counterintuitively, drift can sometimes lead to more rapid loss of a balanced polymorphism than a neutral polymorphism ^84,124,125^, resulting in lower genetic diversity than expected at neutrally evolving loci (Fig. 4B). This outcome is particularly likely when the population-scaled strength of selection is small (*e.g.*, *kN*_*e*_*α* < 10) and equilibrium allele frequencies are close to zero or one ^85,126,127^.

These counterintuitive predictions for balancing selection in finite populations can be viewed as a special case of a broader population genetic phenomenon. Intuition might lead us to predict that loci under balancing selection should exhibit the highest levels of genetic diversity, followed by neutral loci, followed by loci under positive or purifying selection. The classic theory clearly shows that balanced polymorphisms can harbour more or less diversity than neutral loci (Fig. 4B), mirroring results of recent models of genetic diversity for loci under weak positive selection ^128,129^. Thus, there is no one-to-one mapping between balancing selection and inflation of genetic diversity relative to neutral expectations.

#### Multi-locus systems with linkage

The models discussed so far have considered single loci, yet genes can mutually influence each other’s potential for generating balancing selection when they are genetically linked on a chromosome. Most analyses have focused on conditions that maintain polymorphism at a pair of partially linked bi-allelic loci ^88^. In the case of heterozygote advantage, polymorphism can be maintained at linked loci ^87,130^. In trade-off scenarios, linkage tends to expand conditions for the maintenance of polymorphism relative to single-locus models ^131^. For example, a haploid two-locus system can favour a balanced polymorphism in a temporally fluctuating environment ^132,133^, but this is unlikely in otherwise similar single-locus models ^63,113,134^. Conditions for polymorphism also expand under restricted recombination in scenarios of niche and sexual antagonism ^101,135–138^, though the opposite is true for antagonistic pleiotropy ^72^. Finally, loci that are not individually under balancing selection can interact epistatically to maintain a stable two-locus polymorphism, given sufficiently strong linkage between the loci ^139–141^.

Balancing selection can also maintain allelic diversity at neutral sites that are linked to a balanced polymorphism. In cases of heterozygote advantage ^142^, this so-called “associative overdominance” maintains polymorphism when the recombination rate between neutral and selected loci is smaller than the strength of selection ^143^. The diversity at neutral sites increases exponentially with the number of balanced polymorphisms they are linked to ^141^—a prediction that applies to various scenarios, including heterozygote advantage ^86^ and temporally fluctuating selection ^144^. Finally, just as balancing selection can sometimes *reduce* diversity at the selected site relative to neutrality (Fig. 4B), this diversity-reducing effect can extend to linked neutral loci as well ^144^.

#### Traits selected towards an optimum

The models described so far do not explicitly include phenotypic effects; instead, genotypes are assigned fitness values that indirectly reflect selection on phenotypes. However, the lack of a genotype-phenotype map limits the range of questions we can ask about balancing selection. How often should we expect balancing selection to arise among new mutations or segregating genetic variants? Should we expect balanced polymorphisms to be transiently or persistently maintained over time? While the “parameter space” for balancing selection (Box 3) might seem, at first glance, a reasonable proxy for the likelihood of maintaining variation under a given scenario—or a framework for comparing scenarios—it does not differentiate between biologically plausible *vs*. implausible parameter values. We therefore need models that specify the mapping between genotype, phenotype and fitness.

A common way to predict the prevalence of balancing selection is to model the fitness effects of mutations affecting traits selected towards an optimum ^27,36,91^. For example, studies using Fisher’s geometric model ^145^ usually examine trait systems in which adaptive mutations are sufficiently rare that adaptation proceeds by a temporal series of evolutionary steps (“adaptive walks”), with each step corresponding to the invasion of a new adaptive variant ^146^. These mutation-limited dynamics of adaptation are most relevant to trait systems with small mutational targets, or where the mutations’ “scaled” phenotypic effect sizes are large (Fig. 5). Importantly, *scaled* sizes can be large even if *absolute* phenotypic effect sizes of mutations are small, provided the population is near its optimum and/or the number of pleiotropically-associated traits under selection (*i.e.*, trait dimensionality or “complexity”) is high ^38,145,147–149^. Moreover, the very conditions that lead to mutation-limited evolution also lead to heterozygote advantage among the mutations that facilitate the adaptive walk toward the optimum ^36^. An adaptive walk of a diploid population is therefore characterised by a series of short-lived episodes of balancing selection in which individual adaptive alleles are subject to balancing selection immediately following their spread within the population, but these balanced polymorphic states are eventually perturbed by the spread of the next adaptive allele ^36^. These transient dynamics of balancing selection also emerge in trade-off scenarios in which mutations exhibit net heterozygote advantage (Box 3; ^37^).

**Figure 5.**
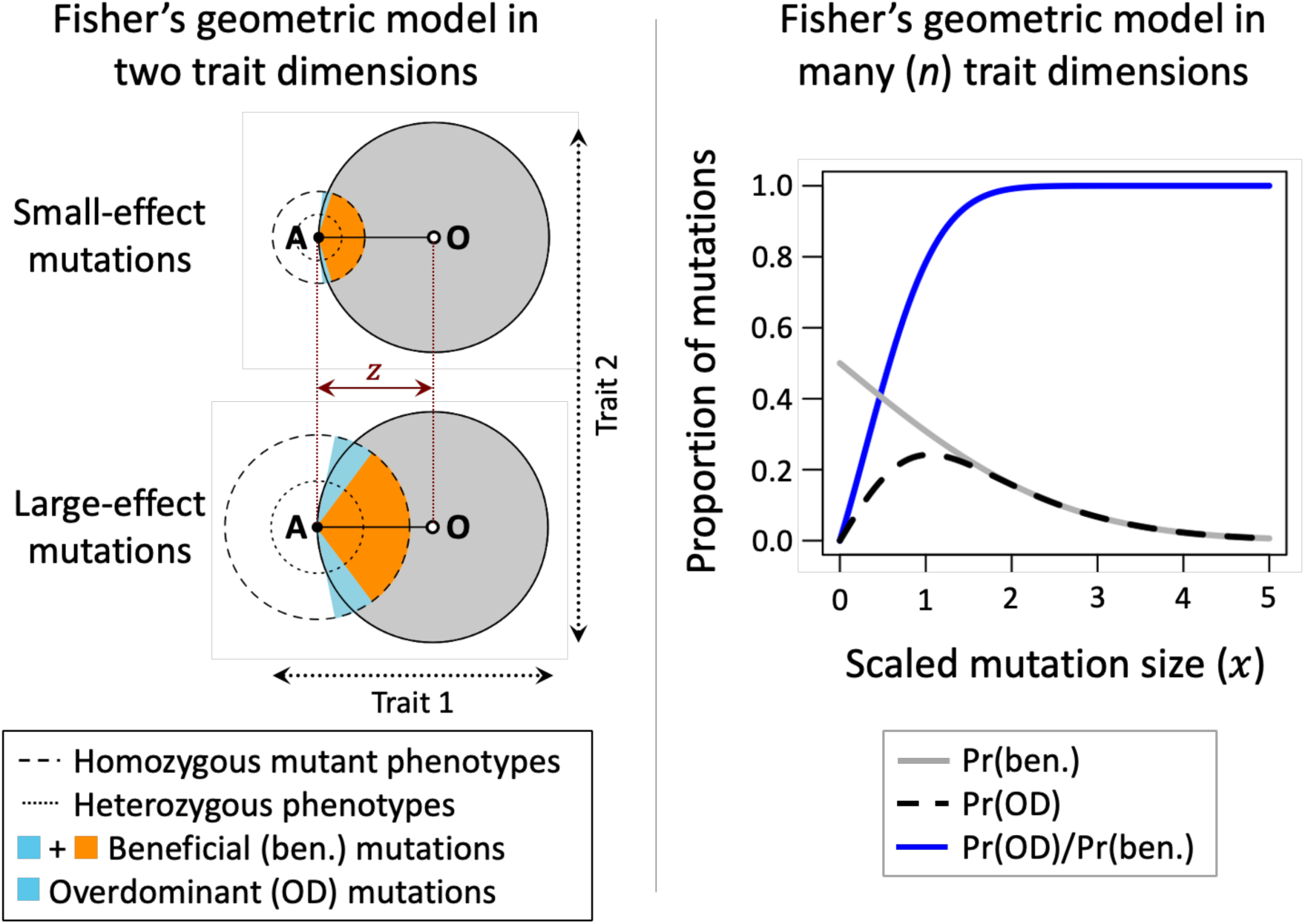
Balancing selection in Fisher’s geometric model. There are *n* traits selected to a single phenotypic optimum defined by the organism’s environment, with **A** denoting the ancestral phenotype (homozygotes for the ancestral allele), **O** representing the optimum phenotype and *z* the distance to the optimum. Fitness declines with the distance between an individual’s phenotype and the optimum, and mutations have random and unbiased orientations in *n*-dimensional phenotypic space. Pleiotropy, which is inherent in the model, can lead to trade-offs, in which mutations causing beneficial changes in some traits simultaneously cause harmful changes in others. The left-hand panel illustrates a series of small-effect mutations (top) and large-effect mutations (bottom) for the case of *n* = 2 traits. Beneficial mutations (where heterozygous carriers are closer to the optimum than ancestral homozygotes) are shaded in orange and blue, with the latter representing mutations under heterozygote advantage. At high dimensions (right-hand panel), the proportion of beneficial variants that exhibit heterozygote advantage is predicted by the scaled mutation size, 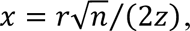 which depends on the number of traits (*n*), and the mutation’s absolute phenotypic effect size (*r*) relative to the distance to the optimum (*z*). “Anisotropy” with respect to selection or mutational effects (*i.e.*, variation among traits in the strength of stabilizing selection, correlational selection, and non-random phenotypic effects of mutations) reduces the *effective* dimensionality of the system ^160–162^.

While transient episodes of balancing selection may occur during the evolutionary approach of a population to an optimum, conditions for long-term balancing selection become restrictive if there is sufficient genetic variation that allows the population to reach its optimum. In such cases, alleles whose homozygous carriers express the optimal phenotype will eventually become fixed (ch. 28 of ^150^), and the ensuing stabilizing selection on the trait will remove rather than preserve genetic variation. If no homozygous genotype corresponds to the optimum, a single long-term balanced polymorphism can be maintained, with the remaining loci experiencing purifying selection against whichever allele is rarest ^26,27^. Still, opportunities for long-term balanced polymorphisms affecting a polygenic trait can expand if loci contributing to the trait are tightly linked ^151,152^, if there is pervasive dominance reversal (Box 3) or “diminishing-returns epistasis” ^43,153^, or if there is strong “disruptive selection” favouring individuals at the trait extremes ^154–156^.

Finally, while stabilizing selection affecting a single quantitative trait tends to remove polymorphism (with the exceptions noted above), balancing selection can still arise when variants underlying the trait have pleiotropic effects on other traits that affect fitness. In these “pleiotropic balancing selection” models ^27,90,157–159^, selection through one fitness component promotes the maintenance of a balanced polymorphism, whereas selection through a second, pleiotropic fitness component—the quantitative trait under stabilizing selection—promotes its removal. Which component predominates depends on the relative strengths of selection through each (see Supplementary Material). Strong balancing selection generated through the first can easily offset comparatively weak stabilizing selection on the second, and vice-versa.

### Linking theories of balancing selection to data

#### Contributions of balancing selection to the maintenance of genetic polymorphisms

There is now ample evidence for balanced polymorphisms in natural populations ^13,31^. For example, “top-down” approaches start with a conspicuous trait polymorphism and then investigate its genetic basis ^163^. A classic example is sickle-cell anaemia, whose prevalence in some populations with histories of malaria exposure is explained by heterozygote advantage maintaining the beta-globin polymorphism ^9^. Top-down approaches can identify recent and potentially ongoing balancing selection, and the ecological context of selection is sometimes known, resulting in rich portraits of the natural history of the balanced polymorphism.

“Bottom-up” approaches, by contrast, look for characteristic genomic signatures of balancing selection ^13,31,164^. Such approaches sidestep the substantial difficulties of measuring fitness in natural populations. They are also trait-agnostic, allowing detection of balancing selection candidates affecting any trait, including those that are difficult to measure with high levels of accuracy or replication. Although bottom-up approaches are best suited for identifying signals of long-term balancing selection (*e.g.*, trans-species polymorphisms, genomic regions with elevated neutral diversity ^31^), experimental evolution can be used to uncover ongoing (and possibly recent) selection, and even to identify the traits that drive these genetic responses. The experimental evolution approach is illustrated by classic studies of *Drosophila* inversion polymorphisms ^12^ and similar recent efforts in other species (*e.g.*, ^165^).

Despite emerging empirical evidence for balancing selection, connections between theory and empirical research remain loose, as a citation network analysis quickly reveals (Fig. 1B,C; see also ^166,167^). Consequently, assessing the prevalence of balancing selection in genomes remains challenging. We don’t know, for instance, whether the fraction of loci influenced by balancing selection is closer to 0.1%, 1%, or 10% (though it is very unlikely to be any higher). Nonetheless, ongoing empirical research to estimate the important parameters of balancing selection models can help guide our thinking. For instance, the theory outlined in ‘Traits selected toward an optimum’ shows that transient balancing selection should be particularly common when evolution proceeds by adaptive walks involving effectively large-effect mutations (recall that mutation “size” is a function of pleiotropy and the distance of the population to the optimum; Fig. 5). Empirical work has indeed revealed many examples of large-effect loci contributing to adaptation of traits with small mutational targets ^163^, including balanced polymorphisms affecting pigmentation, and interactions with pathogens, in *Drosophila* ^168,169^. Moreover, the genetic basis of polygenic traits sometimes includes a mixture of small- and large-effect loci, with the latter possibly maintained under balancing selection. This appears to be the case for cuticular hydrocarbon profiles in *Drosophila* ^170^, age at maturity in Atlantic salmon ^171^, and horn size in Soay sheep ^172^. Large-effect variants at high frequencies are unlikely to be neutral or deleterious, and strongly suggest maintenance by balancing selection ^27^, either due to direct selection on the trait of interest, or indirectly through pleiotropy and selection on another trait.

On the other hand, theory also shows that long-term stable balancing selection is unlikely when trait variation is highly polygenic, selection is stabilizing, and pleiotropy is limited (see section: ‘Traits selected toward an optimum’). In such cases, a trait’s genetic variation is predicted to be attributable to many rare alleles maintained at mutation-selection balance, with up to one locus under balancing selection ^26,27,154^. Such predictions are consistent with the observation that evolutionary responses to artificial selection can occur over short timescales ^173^, and with the absence of intermediate-frequency genetic variants of large-effect in many genome-wide association studies ^174^. Nevertheless, it is important to keep in mind that the populations and traits that have been targeted for intensive study are not necessarily representative or “typical” traits, as each was chosen for personal, societal or logistical reasons. While it is certainly true that many well-studied traits are polygenic and continuously variable, others strongly deviate from the continuous polygenic ideal. In addition to visually conspicuous examples like colour patterns, traits associated with the basic molecular functions of individual genes (*e.g.*, their binding, catalytic, and other cellular functions) have small mutational targets (*e.g.*, genic mutation rates of order 10^-6^) and are likely to display discontinuous patterns of genetic variability. Such “molecular traits” are reasonable candidates for evolution by adaptive walks ^175^, which *are* conducive to short-term episodes of balancing selection ^36,37^. These adaptive walk scenarios are unlikely to generate signals of long-term balancing selection (*e.g.*, inflated linked heterozygosity, gene genealogies with long internal branches, or trans-species polymorphisms), but they can generate signals of short-term balancing selection (*e.g.*, relatively low population genetic differentiation, or partial selective sweeps; ^13,36,164^). Under this view, it is hardly surprising that our most compelling and well-understood examples of balancing selection are both short-term and mediated by selection on fundamental protein structure and function ^9^.

Empirical research is also shedding light on other factors that affect the prevalence of balancing selection. For instance, interactions within and between loci (*e.g.*, dominance reversals; diminishing-returns epistasis) can expand conditions for balancing selection ^43,137,153^. There is evidence for dominance reversals at major loci affecting life-history traits in Atlantic salmon, Soay sheep and *Drosophila* ^171,172,176^, and in studies of quantitative traits such as salinity tolerance in copepods ^177^ and fitness of seed beetles ^178^. Meanwhile, diminishing-returns epistasis for beneficial mutations appears to be common in microbial experiments ^179^. Linkage between loci can also facilitate the establishment and maintenance of multi-locus balanced polymorphisms. Indeed, there are now several examples of large chromosome regions maintained as blocks of differentiated sequences, with evidence for long-term balancing selection ^180^. Examples include chromosomal inversions affecting colour polymorphism in stick insects and damselflies, and life-history traits in seaweed flies ^181–183^. Finally, multi-locus balancing selection becomes likely when selection on traits is disruptive, rather than stabilizing^154–156^. Analyses of large human datasets ^184^, as well as *Drosophila* wing morphology ^185,186^ suggest that stabilizing selection is more common than disruptive selection. However, estimates of nonlinear (*e.g.*, stabilizing and disruptive) selection are notoriously noisy ^29^, and it remains unclear how often disruptive selection might play a role in generating balancing selection.

#### Contributions of balancing selection to genetic variation for fitness and its components

Nucleotide diversity in the *genome* appears to be largely neutral and deleterious. Even so, balancing selection may still contribute substantially to variation in *phenotypes* and *fitness* that have been estimated in several species ^2,3,34,35^. This is because the contribution of a single balanced polymorphism to trait and fitness variance is potentially equivalent to many hundreds of loci at mutation-selection balance ^2,187–189^. For example, the additive genetic variance of a single locus with co-dominant, symmetric and antagonistic effects on two fitness components (*i.e.*, an antagonistic pleiotropy model with parameters *s* = *s*_1_ = *s*_2_ and *h*_1_ = *h*_2_ = 0.5; see Boxes 1 and 3) will be *V*_*A*_ = *s*^2^⁄8, whereas the variance of a locus at mutation-selection balance will be *V*_*A*_ ≈ 2*s*ℎ*μ* (Box 4). Given that mutation rates are typically orders of magnitude smaller than selection coefficients ^190,191^, additive genetic variances due to a single balanced polymorphism should be orders of magnitude greater than variances of a single deleterious polymorphism. Similarly, for polygenic traits where one polymorphism is maintained by balancing selection and the rest are at mutation-selection balance (as predicted by some of the polygenic stabilizing selection models described above), the one balanced locus may account for a large fraction of the trait’s genetic variance, even though most loci affecting the trait evolve under purifying selection.

##### Box 4. Genetic variances due to deleterious and balanced polymorphisms

In randomly mating diploid populations, the contribution of a single biallelic locus to the additive and dominance genetic variance of a trait (*V_A_* and *V_D_*, respectively) is:

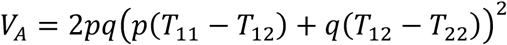

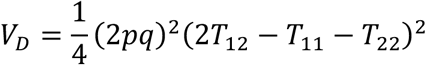

where *p* and *q* are the population frequencies of the *A*_1_ and *A*_2_ alleles, and *T*_11_, *T*_12_, and *T*_22_ represent the average trait (or fitness) values of individuals carrying genotypes *A*_1_*A*_1_, *A*_1_*A*_2_ and *A*_2_*A*_2_, respectively ^34,202^. Thus, the locus’ contribution to each variance component depends on both the proportion of the population that is heterozygous (2*pq*) and the magnitude of genotypic differences in trait values.

Expressions for *V*_*A*_ and *V*_*D*_ can be evaluated under different population genetic models for the maintenance of genetic variation, allowing for comparisons of their predictions. For example, for models of mutation-selection balance, the frequency of a deleterious allele is 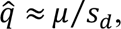 and its contribution to the additive genetic variance for fitness will be *V*_*A*_ ≈ 2*s*_*d*_*μ*, where *μ* is the mutation rate per gamete and *s*_*d*_is the fitness cost to heterozygous carriers of the mutation (the approximations assume that selection is strong relative to mutation: *s*_*d*_ ≫ *μ*). Note that the contribution of deleterious alleles to single fitness components is expected to be even lower than these approximations predict ^2^. For balancing selection models, trade-offs between sexes, niches, or temporal environments will contribute to the additive genetic variance for overall fitness. By contrast, heterozygote advantage and negative frequency-dependent selection contribute no additive genetic variance for overall fitness at equilibrium (Table 2). The same is true for antagonistic pleiotropy (which relies on true heterozygote advantage to maintain the polymorphism; Box 3), though it contributes to additive genetic variance for individual fitness *components* ^2^.

**Table 2.**
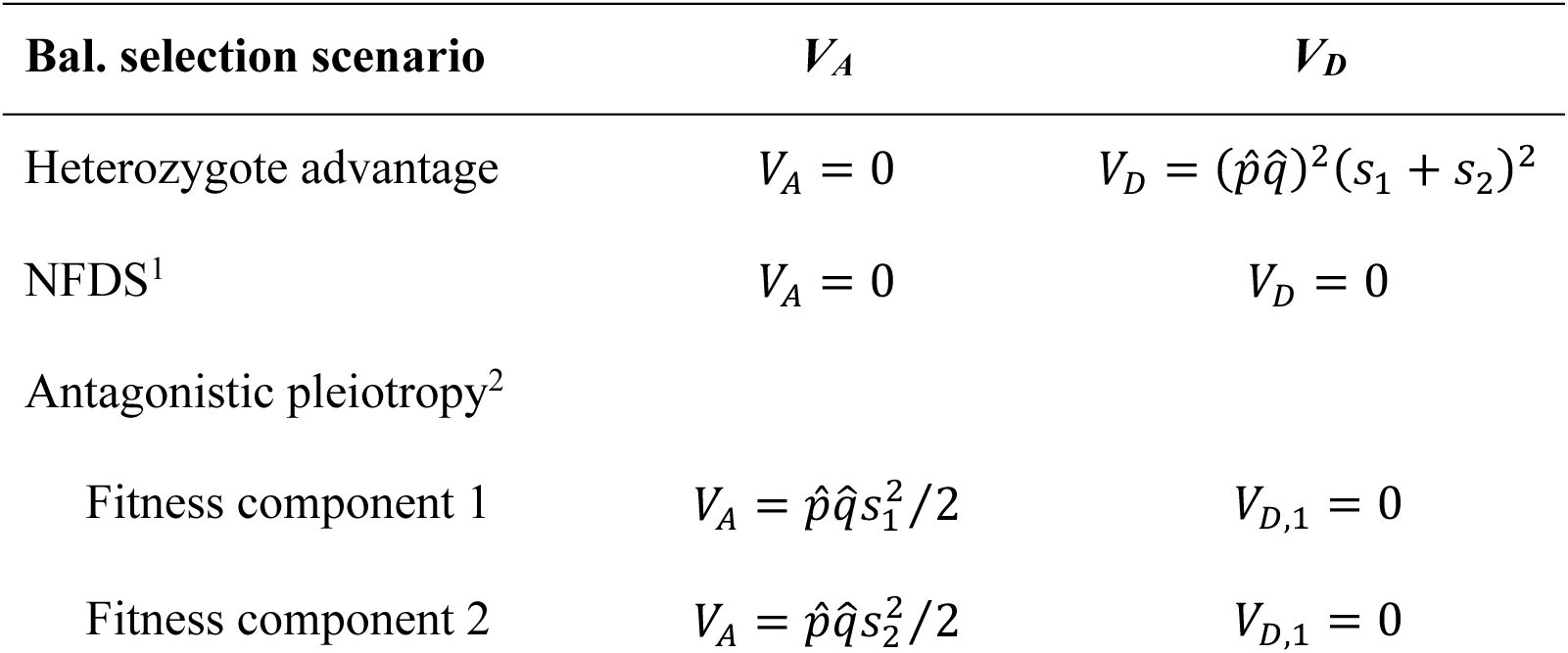
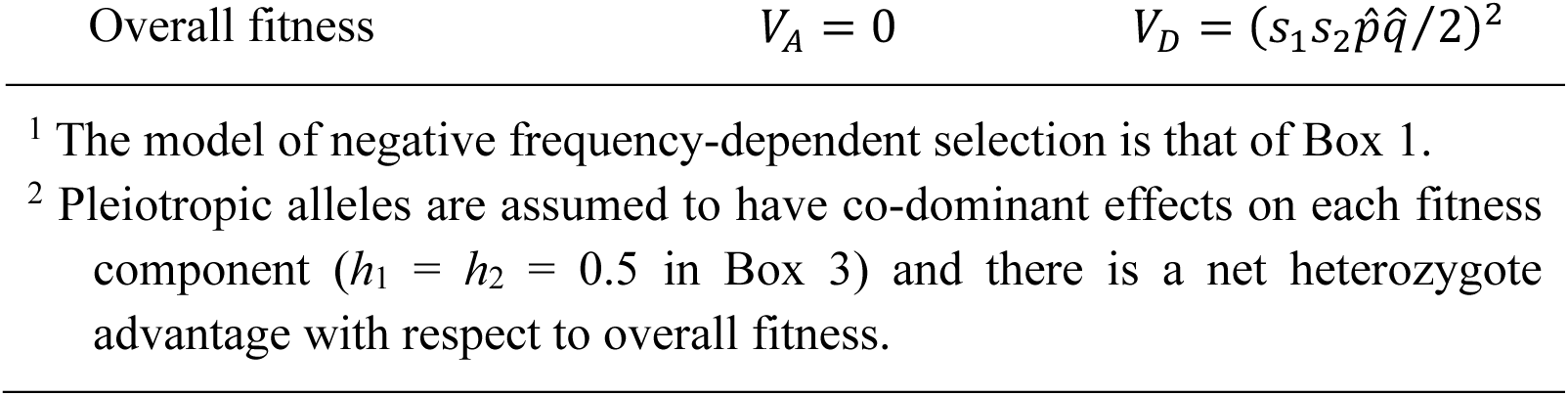
Additive and dominance genetic variance at equilibrium under balancing selection due to heterozygote advantage, negative frequency-dependent selection (NFDS^1^), and a special case of antagonistic pleiotropy^2^.

A final point is that not all balancing selection scenarios maintain *additive* genetic variance for fitness at equilibrium, even though observations of high additive variation for fitness and fitness components are often used as arguments for the prevalence of balancing selection ^2,34,35^. For example, antagonistic pleiotropy, heterozygote advantage and negative frequency-dependent selection do *not* necessarily maintain additive genetic variance for fitness, though they can contribute to the dominance variance for fitness, and the additive variance of individual fitness components (^2^; Box 4). Other trade-off scenarios, including antagonistic selection between niches, sexes, and temporally fluctuating environments, can maintain additive genetic variance for fitness. Thus, while balancing selection might well account for a large fraction of genetic variance for life-history traits and fitness, the different scenarios of balancing selection likely contribute to different components of the genetic variance ^2^. There is, of course, a need for further estimates of the genetic variances of fitness traits, which are currently limited to a small number of populations. Evaluating whether such data consistently exceed what can be explained by mutation-selection balance, and identifying the specific forms of balancing selection that best explain such excesses, remains a pressing empirical challenge.

## Supporting information

Appendix 1 to 4 (Mathematical derivations)

Appendix 5 (Citation analysis)

## Acknowledgements

We thank Brian and Deborah Charlesworth for extensive comments and suggestions. We also thank Göran Arnqvist, Jitka Polechová and Philip Hedrick for further comments. This work was supported by a H2020 Marie Skłodowska-Curie COFUND Action fellowship (#101034413, to FR), the Birgitta Sintring Foundation (#S2024-0007, to MKZ), the Research Council of Norway (302619, to DG), the Alexander von Humboldt Foundation (to HK), the Swiss National Science Foundation (#211549, to XLR), the Swedish Research Council (#2022-03603, to CO; #2020-03123, to EIS) and the European Research Council (ERC-2023-STG-#101117517, to CO). We are particularly grateful to the European Society of Evolutionary Biology for funding a Special Topics Network workshop (to TC, HK, EIS), from which this review began.

