## Appendix 1 to 4 (Mathematical derivations) for "A century of theories of balancing selection"

**Appendix 1. The Basics**

In this appendix we detail the derivation of eq. (1) in the main text under the different scenarios of balancing selection outlined in Box 1 of the main text. Repeating the setup from Section 2 in the main text, we study a diploid, randomly mating population of infinite size, i.e., genetic drift is negligible. Generations are non-overlapping and time is measured in generations. The frequency of allele *A*_1_ is denoted by *p*, and the frequency of the *A*_2_ alleles is *q* = 1 – *p*. The (relative) fitness values of genotypes *A*_1_*A*_1_, *A*_1_*A*_2_ and *A*_2_*A*_2_ in the situations of overdominance, negative frequency dependent selection (NFDS), meiotic drive, and under fitness trade-offs, are defined in Box 1 in the main text.

In eq. (1) in the main text, a canonical form of allele frequency dynamics under balancing selection is stated. It reads:

$$\begin{aligned} \Delta p\approx\alpha p\left( 1-p \right)\left( \hat{p}-p \right) ,\#\left( 1 \right) \end{aligned}$$

where $\Delta p$ denotes the change in allele frequencies from one generation to the next, $\alpha$ describes the effective selection coefficient and $\hat{p}$ is the polymorphic equilibrium state. Balancing selection occurs if $0<\hat{p}<1$ and $\alpha>0$.

Eq. (1) follows from classical population genetic considerations of allele frequency dynamics for a focal allele *A*_1_ at frequency $p\left( t \right)$ at generation *t*. For example, if we define $w_{ij}$ as the fitness associated with genotype *A_i_A_j_*, then the frequency of the *A*_1_ allele in the next generation will be:

$$\begin{aligned} p^{'}=\frac{w_{11}}{\bar{w}}p^{2}+\frac{w_{12}}{\bar{w}}p\left( 1-p \right) ,\#\left( 2 \right) \end{aligned}$$

where $\bar{w}=p^{2}w_{11}+2p\left( 1-p \right)w_{12}+\left( 1-p \right)^{2}w_{22}$ is the average fitness in the population. The change in allele frequency across a single generation is:

$$\begin{aligned} \Delta p=p^{'}-p=\frac{p^{2}w_{11}+p\left( 1-p \right)w_{12}-p\bar{w}}{\bar{w}}\#\left( 3 \right) \end{aligned}$$

$$=\frac{p\left( 1-p \right)\left( w_{12}-w_{22}-p\left( 2w_{12}-w_{11}-w_{22} \right) \right)}{\bar{w}}.$$

If we define $\alpha=2w_{12}-w_{11}-w_{22}$ and $\hat{p}=\left( w_{12}-w_{22} \right)/\left( 2w_{12}-w_{11}-w_{22} \right)$, and assume that the fitness differences between genotypes are small (i.e., selection is weak), then $\bar{w}$ is close to unity, and eq. (3) is well-approximated by eq. (1). We now show how to compute the selection coefficient $\alpha$ and polymorphic equilibrium $\hat{p}$ under each scenario of balancing selection in Box 1 in the main text. Additionally, we provide conditions on the fitness coefficients and dominance parameter(s) for balancing selection to occur ($\alpha>0$ and $0<\hat{p}<1$).

**Overdominance (heterozygote advantage)**

For overdominance, we have $w_{11}=1-s_{1}$, $w_{12}=1$, and $w_{22}=1-s_{2}$, which leads to:

$$\begin{aligned} \Delta p=\frac{p\left( 1-p \right)\left( s_{2}-p\left( s_{1}+s_{2} \right) \right)}{\bar{w}}\approx\left( s_{1}+s_{2} \right)p\left( 1-p \right)\left( \frac{s_{2}}{s_{1}+s_{2}}-p \right) ,\#\left( 4 \right) \end{aligned}$$

where $\alpha=s_{1}+s_{2}$ and $\hat{p}={s_{2}}/\left( s_{1}+s_{2} \right)$, and where the approximation holds in the case of weak selection ($s_{1},s_{2}\ll1$). With overdominance, one always has $s_{1},s_{2}>0$, and the polymorphic equilibrium is, thus, always feasible ($\hat{p}\in\left( 0,1 \right)$) and stable ($\alpha>0$). That is, overdominance induces balancing selection for any parameter value of $0<s_{1}\leq1$ and $0<s_{2}\leq1$ in randomly mating populations.

**Negative frequency-dependent selection (NFDS)**

There are many ways to model frequency-dependent selection, but we consider a model like the one that is presented in Felsenstein (2015). We assume that the fitnesses are frequency-dependent, with $w_{11}=\left( 1-s_{1}p \right)^{2}$, $w_{12}=\left( 1-s_{1}p \right)\left( 1-s_{2} \right)$, and $w_{22}=\left( 1-s_{2} \right)^{2}$. From these, we first compute the average fitness in the population, which is:

$$\begin{aligned} \bar{w}=p^{2}\left( 1-s_{1}p \right)^{2}+2p\left( 1-p \right)\left( 1-s_{1}p \right)\left( 1-s_{2} \right)+\left( 1-p \right)^{2}\left( 1-s_{2} \right)^{2}\#\left( 5 \right) \end{aligned}$$

$$=\left( p\left( 1-s_{1}p \right)+\left( 1-p \right)\left( 1-s_{2} \right) \right)^{2}.$$

Then, from eq. (3), the allele frequency change from one generation to the next is:

$$\begin{aligned} \Delta p=\frac{p\left( 1-p \right)\left( \left( s_{2}-s_{1}p \right)\left( 1-s_{2} \right)+p\left( s_{2}-s_{1}p \right)^{2} \right)}{\bar{w}}\#\left( 6 \right) \end{aligned}$$

$$=\frac{s_{1}p\left( 1-p \right)\left( \frac{s_{2}}{s_{1}}-p \right)}{\sqrt{\bar{w}}}\approx s_{1}p\left( 1-p \right)\left( \frac{s_{2}}{s_{1}}-p \right) ,$$

where $\alpha=s_{1}$ and $\hat{p}={s_{2}}/{s_{1}}$, and where the approximation holds in the case of weak selection ($s_{1},s_{2}\ll1$). The polymorphic equilibrium is feasible ($0<\hat{p}<1$) if $s_{2}<s_{1}$ and always stable if feasible, because $s_{1}=\alpha>0$.

**Meiotic drive**

Drive alleles change the gamete ratio in heterozygotes. Following the expositions in Deredec et al. (2008) and Rode et al. (2019), heterozygotes (on average) produce (1 + *c*)/2 drive alleles, so that *c* represents the deviation from normal Mendelian segregation (*c* = 0 corresponds to fair segregation, and *c* = 1 is the maximum segregation advantage of drive alleles). Letting 1 – *hs* and 1 – *s* represent the heterozygous and homozygous cost of carrying a drive allele (as in Box 1 in the main text), the change in the drive allele frequency across a generation becomes:

$$\begin{aligned} \Delta p=p^{'}-p=\frac{p^{2}\left( 1-s \right)+\left( 1+c \right)p\left( 1-p \right)\left( 1-hs \right)}{\bar{w}}-p\#\left( 7 \right) \end{aligned}$$

$$=\frac{p\left( 1-p \right)\left( c-hs\left( 1+c \right)-ps\left( 1-2h \right) \right)}{\bar{w}}$$

$$\approx s\left( 1-2h \right)p\left( 1-p \right)\left( \frac{c-hs\left( 1+c \right)}{s\left( 1-2h \right)}-p \right) ,$$

where $\alpha=s\left( 1-2h \right)$, $\hat{p}=\left( c-hs\left( 1+c \right) \right)/\left( s\left( 1-2h \right) \right)$, and $\bar{w}=p^{2}\left( 1-s \right)+2p\left( 1-p \right)\left( 1-hs \right)+\left( 1-p \right)^{2}$. The polymorphic equilibrium is feasible ($0<\hat{p}<1$) if:

$$\begin{aligned} \frac{c}{1-h\left( 1-c \right)}<s<\frac{c}{h\left( 1+c \right)} .\#\left( 8 \right) \end{aligned}$$

Balancing selection occurs under meiotic drive if in addition $\alpha=s\left( 1-2h \right)>0$, which is equivalent to *h* < 0.5.

**Antagonistic pleiotropy**

All fitness trade-off scenarios have the same fitness structure (as in Box 1 in the main text), but they vary in the way allele frequency changes are calculated. For example, in the case of antagonistic pleiotropy with each fitness component contributing multiplicatively to overall fitness, the allele frequency changes are given by:

$$\begin{aligned} \Delta p=\frac{p^{2}\left( 1-s_{2} \right)+p\left( 1-p \right)\left( 1-h_{1}s_{1} \right)\left( 1-h_{2}s_{2} \right)}{\bar{w}}-p=\frac{\alpha p\left( 1-p \right)\left( \hat{p}-p \right)}{\bar{w}} ,\#\left( 9 \right) \end{aligned}$$

where $\bar{w}=p^{2}\left( 1-s_{2} \right)+2p\left( 1-p \right)\left( 1-h_{1}s_{1} \right)\left( 1-h_{2}s_{2} \right)+\left( 1-p \right)^{2}\left( 1-s_{1} \right)$ is the mean fitness, $\hat{p}=\left( s_{1}\left( 1-h_{1} \right)-h_{2}s_{2}+s_{1}s_{2}h_{1}h_{2} \right)/\left( s_{1}\left( 1-2h_{1} \right)+s_{2}\left( 1-2h_{2} \right)+2s_{1}s_{2}h_{1}h_{2} \right)$ is the polymorphic equilibrium, and $\alpha=s_{1}\left( 1-2h_{1} \right)+s_{2}\left( 1-2h_{2} \right)+2s_{1}s_{2}h_{1}h_{2}$. Under weak selection ($\bar{w}\approx1$), the expression for allele frequency change simplifies to the canonical form of eq. (1) in the main text.

The condition for balancing selection is:

$$\begin{aligned} \frac{s_{2}h_{2}}{1-h_{1}+s_{2}h_{1}h_{2}}<s_{1}<\frac{s_{2}\left( 1-h_{2} \right)}{h_{1}\left( 1-s_{2}h_{2} \right)} .\#\left( 10 \right) \end{aligned}$$

**Fitness trade-offs between sexes**

With fitness trade-offs between sexes, assume that *A*_1_ is the female-beneficial allele and *A*_2_ is male-beneficial. The female fitness values for genotypes *A*_1_*A*_1_, *A*_1_*A*_2_, and *A*_2_*A*_2_, respectively, are $w_{11,f}=1$, $w_{12,f}=1-s_{1}h_{1}$, and $w_{22,m}=1-s_{1}$. The fitness values for males are $w_{11,m}=1-s_{2}$, $w_{12,m}=1-s_{2}h_{2}$, and $w_{22,m}=1$. Letting *p_f_* and *p_m_* represent the frequency of the *A*_1_ allele in eggs and sperm contributing to fertilization at generation *t*, their values in the next generation will be (Kidwell et al. 1977):

$$\begin{aligned} p_{f}^{'}=\frac{p_{f}p_{m}+\frac{1}{2}\left( p_{f}+p_{m}-2p_{f}p_{m} \right)\left( 1-s_{1}h_{1} \right)}{p_{f}p_{m}+\left( p_{f}+p_{m}-2p_{f}p_{m} \right)\left( 1-s_{1}h_{1} \right)+\left( 1-p_{f} \right)\left( 1-p_{m} \right)\left( 1-s_{1} \right)} ,\#\left( 11a \right) \end{aligned}$$

$$\begin{aligned} p_{m}^{'}=\frac{p_{f}p_{m}\left( 1-s_{2} \right)+\frac{1}{2}\left( p_{f}+p_{m}-2p_{f}p_{m} \right)\left( 1-s_{2}h_{2} \right)}{p_{f}p_{m}\left( 1-s_{2} \right)+\left( p_{f}+p_{m}-2p_{f}p_{m} \right)\left( 1-s_{2}h_{2} \right)+\left( 1-p_{f} \right)\left( 1-p_{m} \right)} .\#\left( 11b \right) \end{aligned}$$

The general condition for balancing selection is:

$$\begin{aligned} \frac{s_{2}h_{2}}{1-h_{1}+s_{2}h_{2}}<s_{1}<\frac{s_{2}\left( 1-h_{2} \right)}{h_{1}\left( 1-s_{2} \right)} .\#\left( 12 \right) \end{aligned}$$

Under weak selection, the allele frequency dynamics can be approximated as:

$$\begin{aligned} p_{f}^{'}\approx\frac{p^{2}+p\left( 1-p \right)\left( 1-s_{1}h_{1} \right)}{p^{2}+2p\left( 1-p \right)\left( 1-s_{1}h_{1} \right)+\left( 1-p \right)^{2}\left( 1-s_{1} \right)} ,\#\left( 13a \right) \end{aligned}$$

$$\begin{aligned} p_{m}^{'}\approx\frac{p^{2}\left( 1-s_{2} \right)+p\left( 1-p \right)\left( 1-s_{2}h_{2} \right)}{p^{2}\left( 1-s_{2} \right)+2p\left( 1-p \right)\left( 1-s_{2}h_{2} \right)+\left( 1-p \right)^{2}} ,\#\left( 13b \right) \end{aligned}$$

which yields the following overall change in frequency:

$$\begin{aligned} \Delta p\approx\frac{s_{1}\left( 1-h_{1}-p\left( 1-2h_{1} \right) \right)p\left( 1-p \right)}{2\left( p^{2}+2p\left( 1-p \right)\left( 1-s_{1}h_{1} \right)+\left( 1-p \right)^{2}\left( 1-s_{1} \right) \right)}\#\left( 14 \right) \end{aligned}$$

$$-\frac{s_{2}\left( h_{2}+p\left( 1-2h_{2} \right) \right)p\left( 1-p \right)}{2\left( p^{2}\left( 1-s_{2} \right)+2p\left( 1-p \right)\left( 1-s_{2}h_{2} \right)+\left( 1-p \right)^{2} \right)} .$$

When fitness effects are codominant ($h_{1}=h_{2}=0.5$), the dynamics simplify to:

$$\begin{aligned} \Delta p\approx\frac{s_{1}s_{2}p\left( 1-p \right)}{2\left( 1-\left( 1-p \right)s_{1} \right)\left( 1-ps_{2} \right)}\left( \frac{s_{1}-s_{2}+s_{1}s_{2}}{2s_{1}s_{2}}-p \right)\#\left( 15 \right) \end{aligned}$$

$$\approx\frac{s_{1}s_{2}}{2}p\left( 1-p \right)\left( \frac{s_{1}-s_{2}+s_{1}s_{2}}{2s_{1}s_{2}}-p \right) .$$

The final approximation matches the canonical form with $\alpha={s_{1}s_{2}}/2$ and $\hat{p}=\left( s_{1}-s_{2}+s_{1}s_{2} \right)/\left( 2s_{1}s_{2} \right)$. With partial-to-complete masking of fitness costs in heterozygotes ($h_{1},h_{2}<0.5$), the dynamics under weak selection can be approximated as:

$$\begin{aligned} \Delta p\approx\frac{p\left( 1-p \right)}{2}\left( s_{1}\left( 1-h_{1}-p\left( 1-2h_{1} \right) \right)-s_{2}\left( h_{2}+p\left( 1-2h_{2} \right) \right) \right)\#\left( 16 \right) \end{aligned}$$

$$=\frac{p\left( 1-p \right)\left( s_{1}\left( 1-2h_{1} \right)+s_{2}\left( 1-2h_{2} \right) \right)}{2}\left( \frac{s_{1}\left( 1-h_{1} \right)-s_{2}h_{2}}{s_{1}\left( 1-2h_{1} \right)+s_{2}\left( 1-2h_{2} \right)}-p \right) .$$

The final expression matches the canonical form with $\alpha=\left( s_{1}\left( 1-2h_{1} \right)+s_{2}\left( 1-2h_{2} \right) \right)/2$ and $\hat{p}=\left( s_{1}\left( 1-h_{1} \right)-s_{2}h_{2} \right)/\left( s_{1}\left( 1-2h_{1} \right)+s_{2}\left( 1-2h_{2} \right) \right)$.

**Fitness trade-offs between niches**

The simplest models of polymorphism maintained due to a trade-off between ecological niches assume that the adults of the population mate randomly after selection, their offspring settle randomly in one of the two niches, and density regulation ensures that the relative proportion of adults emerging from each niche type is constant and independent of the genotype composition of the population (*i.e.*, selection is “soft”; Levene 1953; Christiansen 1975). With exactly two niches, the *A*_1_ allele frequency for adults emerging from niche one is:

$$\begin{aligned} p_{1}=\frac{p^{2}+p\left( 1-p \right)\left( 1-s_{1}h_{1} \right)}{p^{2}+2p\left( 1-p \right)\left( 1-s_{1}h_{1} \right)+\left( 1-p \right)^{2}\left( 1-s_{1} \right)}\#\left( 17 \right) \end{aligned}$$

and the frequency for adults emerging from niche two is:

$$\begin{aligned} p_{2}=\frac{p^{2}\left( 1-s_{2} \right)+p\left( 1-p \right)\left( 1-s_{2}h_{2} \right)}{p^{2}\left( 1-s_{2} \right)+2p\left( 1-p \right)\left( 1-s_{2}h_{2} \right)+\left( 1-p \right)^{2}} .\#\left( 18 \right) \end{aligned}$$

The overall allele frequency change in the metapopulation is then given by the weighted allele frequency changes in the separate niches:

$$\begin{aligned} \Delta p=f\left( p_{1}-p \right)+\left( 1-f \right)\left( p_{2}-p \right) ,\#\left( 19 \right) \end{aligned}$$

where *f* is the proportion of adults emerging from the first niche and 1 – *f* is the proportion emerging from the second. When the two niches produce equal proportions of adults (*f* = 0.5), then the conditions for polymorphism are identical to those of the model of trade-offs between sexes (eq. (12); see Kidwell et al. 1977). Moreover, eq. (19) matches eq. (14), and the same canonical approximations apply for trade-off scenarios between sexes as between niches.

**Parameter space for balancing selection**

Trade-offs are not sufficient conditions for balancing selection. Indeed, only a subset of the possible parameter states of each trade-off model will lead to balancing selection. What, then, is the proportion of the ‘parameter space’ that will lead to balancing selection? For simplicity, we focus on selection scenarios in which fitness costs range from codominant to completely recessive ($0\leq h,h_{1},h_{2}\leq0.5$), as partial or complete dominance of fitness costs (i.e., $0.5<h\leq1$) are usually unamenable to balancing selection. In the models of antagonistic pleiotropy and trade-offs between sexes or niches, we assume for simplicity that dominance is symmetric between the two contexts of selection (*h*_1_ = *h*_2_ = *h*). Cases where *h*_1_, *h*_2_ < 0.5 are usually referred to as “dominance reversals” or “favourable reversals of dominance”, which are known to expand criteria for balancing selection (see Prout 2000; Connallon and Chenoweth 2019; Reid 2022; Grieshop et al. 2024).

Under ***meiotic drive*,** the full parameter space is $0\leq c,s\leq1$, and the condition for balancing selection are described by eq. (8). The proportion of parameter space that generates balancing selection for a given dominance of *h* ($0\leq h\leq0.5$) is:

$$\begin{aligned} f_{bal}=\int_{0}^{h\left( 1-h \right)^{-1}} \frac{c}{h\left( 1+c \right)}dc+1-\frac{h}{1-h}-\int_{0}^{1} \frac{c}{1-h\left( 1-c \right)}dc\#\left( 20 \right) \end{aligned}$$

$$=\frac{\left( 2h-1 \right)\left( h+\log\left( 1-h \right) \right)}{h^{2}} .$$

Under ***antagonistic pleiotropy*,** the parameter space is $0\leq s_{1},s_{2}\leq1$, and the condition for balancing selection is described by eq. (10). The proportion of parameter space that will lead to balancing selection for a given *h* ($0\leq h=h_{1}=h_{2}\leq0.5$) is:

$$\begin{aligned} f_{bal}=1-2\int_{0}^{1} \frac{s_{1}h}{1-h+s_{1}h^{2}}ds_{1}=1-\frac{2h^{2}+2\left( 1-h \right)\log\left( \frac{1-h}{1-h+h^{2}} \right)}{h^{3}} .\#\left( 21 \right) \end{aligned}$$

With ***trade-offs between sexes***, the parameter space is $0\leq s_{1},s_{2}\leq1$, and the condition for balancing selection is described by eq. (12). The proportion of parameter space that will lead to balancing selection for a given *h* ($0\leq h=h_{1}=h_{2}\leq0.5$) is:

$$\begin{aligned} f_{bal}=1-2\int_{0}^{1} \frac{s_{2}h}{1-h+s_{2}h}ds_{1}=1-\frac{2h+2\left( 1-h \right)\log\left( 1-h \right)}{h} .\#\left( 22 \right) \end{aligned}$$

Eq. (22) applies to the simple model of trade-offs between two equally abundant niches.

**Appendix 2. Balancing selection in haploids and haplodiploids**

**Fitness in haploid and diploid individuals**

Following the baseline models from Appendix 1, we track the evolutionary dynamics of two alleles, *A*_1_ and *A*_2_, which occur at frequencies *p* and *q*, respectively. We assume that the fitness of an individual that is haploid for a given allele is equivalent to that of a diploid individual that is homozygous for the same allele. Table S1 summarizes the haploid and diploid fitness values of each genotype under each model of selection.

| **Table S1.** Fitness of haploid (*A*_1_ and *A*_2_) and diploid genotypes (*A*_1_*A*_1_, *A*_1_*A*_2_, *A*_2_*A*_2_) | | | |
| --- | --- | --- | --- |
|  |  | **Genotype** |  |
|  | ***A*_1_*A*_1_ or *A*_1_** | ***A*_1_*A*_2_** | ***A*_2_*A*_2_ or *A*_2_** |
| **Overdominance:** | 1 – *s*_1_ | 1 | 1 – *s*_2_ |
| **NFDS:** | (1 – *s*_1_*p*)^2^ | (1 – *s*_1_*p*)(1 – *s*_2_) | (1 – *s*_2_)^2^ |
| **Meiotic drive** | 1 | 1 – *sh* | 1 – *s* |
| **Antagonistic selection** |  |  |  |
| **Fitness context 1:** | 1 | 1 – *h*_1_*s*_1_ | 1 – *s*_1_ |
| **Fitness context 2:** | 1 – *s*_2_ | 1 – *h*_2_*s*_2_ | 1 |

**Balancing selection in entirely haploid populations**

Models of overdominance, meiotic drive, and antagonistic pleiotropy cannot maintain a balanced polymorphism in a haploid population. However, the other scenarios of selection from Table S1 can maintain polymorphism. For example, in the ***negative frequency-dependent selection*** (NFDS) model presented in Table S1, the conditions for polymorphism are just as permissive in haploids as they are in diploids.

When there is a ***fitness trade-off between the sexes***, with *A*_1_ as the female-beneficial allele and *A*_2_ as the male-beneficial allele, the conditions for polymorphism can be shown to be the same as those predicted under a diploid model with codominant fitness effects within each sex. At birth, the frequency of individuals with the *A*_1_ genotype is *p*, and the frequency with the *A*_2_ genotype is *q* = 1 – *p*. After selection, the *A*_1_ frequency in females and males, respectively, will be:

$$\begin{aligned} p_{f}=\frac{p}{p+q\left( 1-s_{1} \right)} ,\#\left( 23a \right) \end{aligned}$$

$$\begin{aligned} p_{m}=\frac{p\left( 1-s_{2} \right)}{p\left( 1-s_{2} \right)+q} .\#\left( 23b \right) \end{aligned}$$

Because each sex makes an equal genetic contribution to the next generation, the allele frequency at birth in the next generation will be $p^{'}=\frac{1}{2}\left( p_{f}+p_{m} \right)$, and the change in frequency across a single generation is:

$$\begin{aligned} \Delta p=p^{'}-p=\frac{s_{1}s_{2}p\left( 1-p \right)\left( \hat{p}-p \right)}{\left( 1-ps_{2} \right)\left( 1-qs_{1} \right)}\approx s_{1}s_{2}p\left( 1-p \right)\left( \hat{p}-p \right) ,\#\left( 24 \right) \end{aligned}$$

with the final approximation (appropriate under weak selection) matching the canonical form with $\alpha=s_{1}s_{2}$ and $\hat{p}=\left( s_{1}-s_{2}+s_{1}s_{2} \right)/\left( 2s_{1}s_{2} \right)$. The condition for balancing selection is:

$$\begin{aligned} \frac{s_{2}}{1+s_{2}}<s_{1}<\frac{s_{2}}{1-s_{2}} ,\#\left( 25 \right) \end{aligned}$$

which matches the special case of eq. (12) where $h_{1}=h_{2}=0.5$.

In the haploid model of a ***trade-off between niches*** assume (as in Appendix 1) that there are two habitat types that are distributed randomly across a large number of subpopulations of the species. Individuals experience viability selection in the subpopulation in which they settle as juveniles. The surviving adults emerge from their subpopulation and randomly mate and reproduce offspring of the next generation. These offspring settle at random among subpopulations, which homogenizes the subpopulations with respect to juvenile genotypes. In other words, dispersal and gene flow is unrestricted among subpopulations, which as Levene (1953) notes defines a worst-case-scenario for the maintenance of stable polymorphism.

Let *c_i_* be the fraction of subpopulations that are of the *i*^th^ type (*i* = {1, 2}). There is viability selection in each subpopulation, followed by random mating of adults, whose offspring make up the next generation. With this life cycle, the genotype frequencies will be the same in each habitat prior to selection. The *A*_1_ genotype frequencies after viability selection will be:

$$\begin{aligned} p_{1}=\frac{p}{1-qs_{1}}\#\left( 26a \right) \end{aligned}$$

in habitat 1, and:

$$\begin{aligned} p_{2}=\frac{p\left( 1-s_{2} \right)}{1-ps_{2}}\#\left( 26b \right) \end{aligned}$$

in habitat 2.

Under ‘soft selection’, which follows Levene (1953), a fixed fraction of individuals emerges from each habitat (*i.e.*, selection is “soft”; Christiansen 1975). Supposing the fraction of individuals emerging from each habitat (*c*_1_ and *c*_2_) is proportional to the relative abundance of the two habitats, then the frequency of *A*_1_ among the pool of individuals contributing to reproduction will be $c_{1}p_{1}+c_{2}p_{2}$. The *A*_1_ genotype frequency in the next generation will be $p^{'}=c_{1}p_{1}+c_{2}p_{2}$, and the change in genotype frequency will be:

$$\begin{aligned} \Delta p=c_{1}\left( p_{1}-p \right)+c_{2}\left( p_{2}-p \right)=\frac{s_{1}s_{2}p\left( 1-p \right)\left( \hat{p}-p \right)}{\left( 1-ps_{2} \right)\left( 1-qs_{1} \right)} ,\#\left( 27 \right) \end{aligned}$$

where $\hat{p}=\left( c_{1}s_{1}-c_{2}s_{2}+c_{1}s_{1}s_{2} \right)/\left( s_{1}s_{2} \right)$. Under weak selection, $\Delta p$ simplifies to the canonical form with $\alpha=s_{1}s_{2}$. The conditions for polymorphism are:

$$\begin{aligned} \frac{c_{2}s_{2}}{c_{1}\left( 1+s_{2} \right)}<s_{1}<\frac{c_{2}s_{2}}{c_{1}-s_{2}\left( 1-c_{1} \right)} ,\#\left( 28 \right) \end{aligned}$$

which match those of the model of trade-offs between sexes (eq. (25)) when $c_{1}=c_{2}=0.5$.

Under ‘hard selection’, following Christiansen (1975) and Dempster (1955), the contribution of each habitat type to the pool of reproducing adults depends on both the relative proportions of each habitat (*c*_1_ and *c*_2_) and the mean viability of individuals in each habitat. Taking both factors into account, the frequency of *A*_1_ among the pool of individuals contributing to reproduction will be:

$$\begin{aligned} \frac{c_{1}\left( 1-qs_{1} \right)}{c_{1}\left( 1-qs_{1} \right)+c_{2}\left( 1-ps_{2} \right)}p_{1}+\frac{c_{2}\left( 1-ps_{2} \right)}{c_{1}\left( 1-qs_{1} \right)+c_{2}\left( 1-ps_{2} \right)}p_{2} .\#\left( 29 \right) \end{aligned}$$

The change in frequency of the *A*_1_ allele across a generation is:

$$\begin{aligned} \Delta p=\frac{\left( c_{1}s_{1}-c_{2}s_{2} \right)p\left( 1-p \right)}{c_{1}\left( 1-qs_{1} \right)+c_{2}\left( 1-ps_{2} \right)}\#\left( 30 \right) \end{aligned}$$

and there no possibility of a polymorphic equilibrium. Rather, the genotype with the highest arithmetic mean fitness (accounting for the relative abundances of the two habitats) is expected to become fixed and the other genotype eliminated by selection.

While hard selection reduces scope for maintaining polymorphism, two other factors can expand the scope for maintaining polymorphism. First, factors that promote habitat differentiation of genotype frequencies can expand the conditions for polymorphism. Examples include preferential dispersal of individuals into habitats to which they are better adapted (as might arise due to divergent habitat preferences of different genotypes), and limited dispersal between habitats, which leads to preferential mating and reproduction among individuals from the same versus different subpopulations. Second, diploidy can greatly improve prospects for polymorphism when locally adapted alleles tend to be relatively dominant (i.e., *h*_1_, *h*_2_ < 0.5), which leads to masking of the fitness costs of locally maladapted variants. The core theory in diploids is described in Levene (1953), Dempster (1955), and Christiansen (1975). Extensions to haploid populations can be found in Gliddon and Strobeck (1975) and Czochor and Leonard (1982). The importance of dominance reversals in diploids was stressed by Hoekstra, Bijlsma and Dolman (1985).

**Balancing selection in haplo-diploid populations and at X-linked loci**

We focus on haplo-diploid species in which the female is diploid and the male is haploid. These models also apply to X-linked genomic regions in species with a degenerate Y chromosome. The models are also relevant to Z-linked genes in species with a degenerate W chromosome, provided the sex labels are reversed (females become haploid; males become diploid). Allele frequencies typically diverge between sexes for all models of selection, which makes it convenient to track allele frequencies in the female and male gametes that contribute to the next generation. Let *p_f_* and *p_m_* represent the frequency of the *A*_1_ allele in the eggs and sperm (respectively) that contribute to fertilization of the females of the next generation. Since males inherit an X chromosome from their mothers, *p_f_* will be the frequency with which males of the next generation inherit an *A*_1_ allele. We focus on models that generate qualitatively different outcomes relative to diploid models. Hence, we ignore NFDS beyond noting that the model outlined in Table S1 is sufficient for maintaining an X-linked polymorphism.

Under weak selection and specific conditions of dominance (which we specify below), the evolutionary dynamics of haplodiploid and X-linked loci are well-approximated as $\Delta p\approx\frac{2}{3}\Delta p_{f}+\frac{1}{3}\Delta p_{m}$, where $\Delta p_{f}$ and $\Delta p_{m}$ represent the within-generation response to selection in females and males, respectively. To first order in the selection coefficients (*i.e.*, the differences in fitness between genotypes), the within generation responses are:

$$\begin{aligned} \Delta p_{f}\approx p\left( 1-p \right)\left( p\left( w_{11}-w_{12} \right)+\left( 1-p \right)\left( w_{12}-w_{22} \right) \right) ,\#\left( 31a \right) \end{aligned}$$

$$\begin{aligned} \Delta p_{m}\approx p\left( 1-p \right)\left( v_{1}-v_{2} \right) ,\#\left( 31b \right) \end{aligned}$$

where *w*_11_, *w*_12_, and *w*_22_ represent the female fitness values associated with genotypes *A*_1_*A*_1_, *A*_1_*A*_2_, and *A*_2_*A*_2_, respectively, and *v*_1_ and *v*_2_ are the male fitness values for genotypes *A*_1_ and *A*_2_.

With ***overdominant selection***, the exact allele frequency changes in female and male gametes are:

$$\begin{aligned} \Delta p_{f}=\frac{p_{f}p_{m}\left( 1-s_{1} \right)+\frac{1}{2}\left( p_{f}+p_{m}-2p_{f}p_{m} \right)}{p_{f}p_{m}\left( 1-s_{1} \right)+\left( p_{f}+p_{m}-2p_{f}p_{m} \right)+q_{f}q_{m}\left( 1-s_{2} \right)}-p_{f} ,\#\left( 32a \right) \end{aligned}$$

$$\begin{aligned} \Delta p_{m}=\frac{p_{f}\left( 1-s_{1} \right)}{p_{f}\left( 1-s_{1} \right)+q_{f}\left( 1-s_{2} \right)}-p_{m} ,\#\left( 32b \right) \end{aligned}$$

which yields the exact polymorphic equilibrium frequencies:

$$\begin{aligned} \hat{p}_{f}=\frac{s_{2}\left( \frac{3}{2}-s_{2} \right)-\frac{1}{2}s_{1}}{s_{1}\left( 1-s_{1} \right)+s_{2}\left( 1-s_{2} \right)} ,\#\left( 33a \right) \end{aligned}$$

$$\begin{aligned} \hat{p}_{m}=\frac{\hat{p}_{f}\left( 1-s_{1} \right)}{1-\hat{p}_{f}s_{1}-\hat{q}_{f}s_{2}} .\#\left( 33b \right) \end{aligned}$$

Polymorphism occurs when the following condition is true (see Pamilo 1979):

$$\begin{aligned} \frac{3}{4}-\frac{3}{4}\sqrt{1-\frac{8}{9}s_{1}}<s_{2}<s_{1}\left( 3-2s_{1} \right) .\#\left( 34 \right) \end{aligned}$$

Under weak selection (*s*_1_, *s*_2_ << 1), the allele frequency dynamics can be approximated by:

$$\begin{aligned} \Delta p\approx\frac{2}{3}\left( s_{1}+s_{2} \right)p\left( 1-p \right)\left( \hat{p}-p \right) ,\#\left( 35 \right) \end{aligned}$$

which matches the canonical form with $\alpha=\frac{2}{3}\left( s_{1}+s_{2} \right)$ and $\hat{p}=\left( 3s_{2}-s_{1} \right)/\left( 2\left( s_{1}+s_{2} \right) \right)$. The (approximate) condition for maintaining polymorphic becomes:

$$\begin{aligned} \frac{1}{3}s_{1}<s_{2}<3s_{1} .\#\left( 36 \right) \end{aligned}$$

With ***X-linked meiotic drive in females and fitness costs in both sexes*** (*i.e.*, meiotic drive occurs in females and costs of the drive allele are expressed by both sexes), the exact allele frequency dynamics of the driving allele will be:

$$\begin{aligned} \Delta p_{f}=\frac{p_{f}p_{m}\left( 1-s \right)+\frac{1+c}{2}\left( p_{f}+p_{m}-2p_{f}p_{m} \right)\left( 1-sh \right)}{p_{f}p_{m}\left( 1-s \right)+\left( p_{f}+p_{m}-2p_{f}p_{m} \right)\left( 1-sh \right)+q_{f}q_{m}}-p_{f} ,\#\left( 37a \right) \end{aligned}$$

$$\begin{aligned} \Delta p_{m}=\frac{p_{f}\left( 1-s \right)}{p_{f}\left( 1-s \right)+q_{f}}-p_{m} .\#\left( 37b \right) \end{aligned}$$

The polymorphic equilibrium (when it exists) corresponds to the allele frequencies:

$$\begin{aligned} \hat{p}_{f}=\frac{2c-s\left( 1+2h-sh \right)\left( 1+c \right)}{2s\left( 1-2h-s\left( 1-h \right) \right)} ,\#\left( 38a \right) \end{aligned}$$

$$\begin{aligned} \hat{p}_{m}=\frac{\hat{p}_{f}\left( 1-s \right)}{1-\hat{p}_{f}s} .\#\left( 38b \right) \end{aligned}$$

Assuming weak selection, weak drive and at least partial recessivity of the cost of the drive allele (*s*, *c* << 1; *h* < ½), we obtain the approximation:

$$\begin{aligned} \Delta p\approx\frac{2}{3}s\left( 1-2h \right)p\left( 1-p \right)\left( \hat{p}-p \right) ,\#\left( 39 \right) \end{aligned}$$

which, given $\alpha=\frac{2}{3}s\left( 1-2h \right)$ and $\hat{p}=\left( 2c-s\left( 1+2h \right) \right)/\left( 2s\left( 1-2h \right) \right)$, matches the canonical form.

X-linked segregation distortion in males (***sex-ratio drive***), is more complex, leads to sex ratio biases in the population, and has received considerable theoretical and empirical attention. We refer readers to Edwards (1961) and Mackintosh et al. (2021) for the classic treatment and recent advances.

***Antagonistic pleiotropy.*** Under antagonistic pleiotropy with fitness components having multiplicative effects on overall fitness, the fitness of *A*_1_*A*_1_ and *A*_1_ individuals is 1 – *s*_1_, the fitness of *A*_2_*A*_2_ and *A*_2_ individuals is 1 – *s*_2_, and fitness of *A*_1_*A*_2_ individuals is (1 – *s*_1_*h*)(1 – *s*_2_*h*), where *h* = *h*_1_ = *h*_2_ is the degree to which the maladaptive effects of each locus are masked in heterozygotes. The exact allele frequency dynamics are:

$$\begin{aligned} \Delta p_{f}=\frac{p_{f}p_{m}\left( 1-s_{1} \right)+\frac{1}{2}\left( p_{f}+p_{m}-2p_{f}p_{m} \right)\left( 1-s_{1}h \right)\left( 1-s_{2}h \right)}{p_{f}p_{m}\left( 1-s_{1} \right)+\left( p_{f}+p_{m}-2p_{f}p_{m} \right)\left( 1-s_{1}h \right)\left( 1-s_{2}h \right)+q_{f}q_{m}\left( 1-s_{2} \right)}-p_{f}\#,\left( 40a \right) \end{aligned}$$

$$\begin{aligned} \Delta p_{m}=\frac{p_{f}\left( 1-s_{1} \right)}{p_{f}\left( 1-s_{1} \right)+q_{f}\left( 1-s_{2} \right)}-p_{m} .\#\left( 40b \right) \end{aligned}$$

Under weak selection with partial masking of maladaptive effects (*s*_1_, *s*_2_ << 1 and *h* < 0.5), the allele frequency dynamics can be approximated by:

$$\begin{aligned} \Delta p\approx\frac{2}{3}\left( s_{1}+s_{2} \right)\left( 1-2h \right)p\left( 1-p \right)\left( \hat{p}-p \right) ,\#\left( 41 \right) \end{aligned}$$

where $\alpha=\frac{2}{3}\left( s_{1}+s_{2} \right)\left( 1-2h \right)$ and $\hat{p}=\left( s_{2}\left( 3-2h \right)-s_{1}\left( 1+2h \right) \right)/\left( 2\left( s_{1}+s_{2} \right)\left( 1-2h \right) \right)$. The (approximate) condition for balanced polymorphism becomes:

$$\begin{aligned} \frac{s_{1}\left( 1+2h \right)}{3-2h}<s_{2}<\frac{s_{1}\left( 3-2h \right)}{1+2h} ,\#\left( 42 \right) \end{aligned}$$

which simplifies to the standard overdominance model when masking is complete (*h* = 0).

When there is a ***trade-off between the sexes***, with the *A*_1_ allele favoured in females and the *A*_2_ allele favoured in males (*i.e.*, $s_{1}=s_{f}$, $s_{2}=s_{m}$, $h_{1}=h_{f}$, $h_{2}=h_{m}$), the allele frequency dynamics are described by:

$$\begin{aligned} \Delta p_{f}=\frac{p_{f}p_{m}+\frac{1}{2}\left( p_{f}+p_{m}-2p_{f}p_{m} \right)\left( 1-s_{f}h_{f} \right)}{p_{f}p_{m}+\left( p_{f}+p_{m}-2p_{f}p_{m} \right)\left( 1-s_{f}h_{f} \right)+q_{f}q_{m}\left( 1-s_{f} \right)}-p_{f} ,\#\left( 43a \right) \end{aligned}$$

$$\begin{aligned} \Delta p_{m}=\frac{p_{f}\left( 1-s_{m} \right)}{p_{f}\left( 1-s_{m} \right)+q_{f}}-p_{m} .\#\left( 43b \right) \end{aligned}$$

The polymorphic equilibrium (when it exists) corresponds to the allele frequencies (see Haldane and Jayakar 1964; Patten and Haig 2009):

$$\begin{aligned} \hat{p}_{f}=\frac{2s_{f}\left( 1-h_{f} \right)-s_{m}+s_{f}s_{m}h_{f}}{2s_{f}\left( 1-2h_{f}+s_{m}h_{f} \right)} ,\#\left( 44a \right) \end{aligned}$$

$$\begin{aligned} \hat{p}_{m}=\frac{\hat{p}_{f}\left( 1-s_{m} \right)}{\hat{p}_{f}\left( 1-s_{m} \right)+\hat{q}_{f}} .\#\left( 44b \right) \end{aligned}$$

The condition for a balanced polymorphism is:

$$\begin{aligned} \frac{2s_{f}h_{f}}{1+s_{f}h_{f}}<s_{m}<\frac{2s_{f}\left( 1-h_{f} \right)}{1-s_{f}h_{f}} .\#\left( 45 \right) \end{aligned}$$

It has been noted that the condition for polymorphism on the X is identical to that of the autosomes when $h_{m}=\left( 2-s_{m} \right)^{-1}$, the conditions are broader on the X than the autosomes when $h_{m}>\left( 2-s_{m} \right)^{-1}$, and they are broader on the autosomes than the X when $h_{m}<\left( 2-s_{m} \right)^{-1}$ (Ruzicka and Connallon 2020).

Under weak selection and at least partial masking of female-deleterious effects in females (*s*_1_, *s*_2_ << 1, and *h_f_* < 0.5), the allele frequency dynamics can be approximated as:

$$\begin{aligned} \Delta p\approx\frac{2}{3}s_{f}\left( 1-2h_{f} \right)p\left( 1-p \right)\left( \hat{p}-p \right)\#\left( 46 \right) \end{aligned}$$

where $\alpha=\frac{2}{3}s_{f}\left( 1-2h_{f} \right)$ and $\hat{p}=\left( 2s_{f}\left( 1-h_{f} \right)-s_{m} \right)/\left( 2s_{f}\left( 1-2h_{f} \right) \right)$ (Albert and Otto 2005; Connallon and Clark 2011, 2013).

Other cases of interest for the polymorphism maintained in haplodiploid populations or X-linked loci include ***multiple-niche polymorphism,*** as addressed by Moody (1979), who showed that conditions for polymorphism were restricted on the X relative to the autosomes. Immler et al. (2012) considered the effects of ***ploidally antagonistic selection*** involving trade-offs between the haploid and diploid phases of a life cycle, including at autosomal and X-linked genes.

**Appendix 3. Balancing selection in finite populations**

The amount of genetic diversity in finite populations reflects a balance between mutation, selection, and genetic drift. The stationary distribution for allele frequencies under mutation, selection and drift provides predictions about the probabilities of observing various allele frequency states in populations at the long-term equilibrium for these processes. Following Wright (1945) and Robertson (1962), the general form of the stationary distribution for a bi-allelic locus is:

$$\begin{aligned} f\left( p \right)=\frac{C}{p\left( 1-p \right)}\exp\left( 2\int\frac{M}{V}dp \right) ,\#\left( 47 \right) \end{aligned}$$

where $f\left( p \right)$ is the probability density function for the *A*_1_ allele, *C* is a constant that ensures that $\int_{0}^{1} f\left( p \right)dp=1$, and *M* and *V* describe the expected value and the variance of allele frequency change across a single generation. With mutation and balancing selection, the expected change in frequency given *p* is:

$$\begin{aligned} M=\alpha p\left( 1-p \right)\left( \hat{p}-p \right)+u\left( 1-p \right)-vp ,\#\left( 48 \right) \end{aligned}$$

where $\alpha$ and $\hat{p}$ are as described in Appendix 1 and 2, $u$ is the rate at which an *A*_2_ allele mutates to an *A*_1_ allele, and $v$ is the rate at which an *A*_1_ allele mutates to an *A*_2_ allele. The variance of allele frequency change (under a standard Wright-Fisher model) is:

$$\begin{aligned} V=\frac{p\left( 1-p \right)}{kN_{e}} ,\#\left( 49 \right) \end{aligned}$$

where *N_e_* is the effective population size and *k* is a constant that reflects the ploidy level of the population or locus in question (*k* = 1 for haploid populations, *k* = 2 for diploids, and *k* = 1.5 in haplo-diploid populations where drift is equally strong through females and males). Substituting eqs. (48) and (49) into eq. (47) and carrying out the integration, and some algebra, yields:

$$\begin{aligned} f\left( p \right)=cp^{2kN_{e}u-1}\left( 1-p \right)^{2kN_{e}v-1}\exp\left( -kN_{e}\alpha\left( \hat{p}-p \right)^{2} \right) ,\#\left( 50 \right) \end{aligned}$$

where *c* ensures that the function integrates to one.

**Appendix 4. Effects of balanced polymorphisms on the expression of traits selected to an optimum**

**Balancing selection in Fisher’s geometric model**

Fisher’s geometric model is arguably the simplest model of random mutation and selection of a set of phenotypic traits. In the isotropic version of Fisher’s model, which we focus on here, there are *n* trait axes, each selected to an optimum. Mutations are randomly oriented in *n*-dimensional trait space, and fitness of individuals is a Gaussian function of the Euclidean distance between their phenotypes and the location of the *n*-dimensional optimum. We will also assume that mutations have codominant effects on trait expression. However, the nonlinear mapping between phenotype and fitness leads to dominance with respect to the fitness effects of the mutations. The population is initially fixed for a wild-type genotype that is displaced from the optimum by amount *z*. Mutation size is quantified using Fisher’s scaling, $x={r\sqrt{n}}/{2z}$ (see Orr 1998), which is a function of the raw phenotypic effect size of the mutation (*r* in Euclidean distance), the distance of wild-type individuals from the optimum (*z*), and the total number of trait dimensions (*n*). Additional complexity arising from differences in selection among developmental stages, environmental variation, and sex differences in selection can be integrated into these models by rescaling mutation size to incorporate what effectively amounts to the greater dimensionality inherent in these more complex selection scenarios (Connallon and Clark 2014).

Fisher (1930) originally used the model to address the probability that a random mutation was beneficial, and showed that this probability was a declining function of the scaled mutation size, *x*. Indeed, the probability that a mutation with phenotypic effect size *x* is beneficial and favoured to increase in frequency within the population (at least when rare) is approximately:

$$\begin{aligned} \Pr\left( \text{ben.}|x \right)=\frac{1}{2}-\frac{1}{2}\mathrm{erf} \left( \frac{x}{2\sqrt{2}} \right) ,\#\left( 51 \right) \end{aligned}$$

where erf() refers to the error function. This expression is accurate when the number of trait dimensions under selection is at least reasonably large (*n* > ~10). Early versions of Fisher’s model either focused on haploid systems or did not explicitly consider the effects of diploidy on the evolutionary dynamics of mutations.

Sellis et al. (2011) and Manna et al. (2011) were the first to emphasize that Fisher’s model, when applied to diploid populations, predicts that beneficial mutations (those that are favoured to invade when rare) fall into two categories. They can experience positive selection, in which they are favoured to sweep to fixation, or they can exhibit heterozygote advantage when paired with the ancestral allele, which favours their evolution to a transient balanced polymorphic state that persists until the next beneficial mutation to spread perturbs the system and leads to a new transient equilibrium state. In a population initially fixed for an ancestral allele, the probability that a new mutation has a heterozygote advantage and is, therefore, subject to transient balancing selection is (Sellis et al. 2011; Connallon and Clark 2014):

$$\begin{aligned} \Pr\left( \text{bal.}|x \right)=\frac{1}{2}\mathrm{erf} \left( \frac{3x}{2\sqrt{2}} \right)-\frac{1}{2}\mathrm{erf} \left( \frac{x}{2\sqrt{2}} \right) .\#\left( 52 \right) \end{aligned}$$

Among the set beneficial mutations with phenotypic effect size *x*, the proportion subject to balancing selection is:

$$\begin{aligned} \frac{\Pr\left( \text{bal.}|x \right)}{\Pr\left( \text{ben.}|x \right)}=\frac{\mathrm{erf} \left( \frac{3x}{2\sqrt{2}} \right)-\mathrm{erf} \left( \frac{x}{2\sqrt{2}} \right)}{1-\mathrm{erf} \left( \frac{x}{2\sqrt{2}} \right)} .\#\left( 53 \right) \end{aligned}$$

The following figure shows the proportions of random mutations that are beneficial ($\Pr\left( \text{ben.}|x \right)$; grey curve) and subject to short-term balancing selection ($\Pr\left( \text{bal.}|x \right)$; black dashed curve) as a function of the scaled mutation size, *x*. The blue curve shows the ratio of the two expressions (i.e., the proportion of beneficial mutations that is subject to short-term balancing selection).

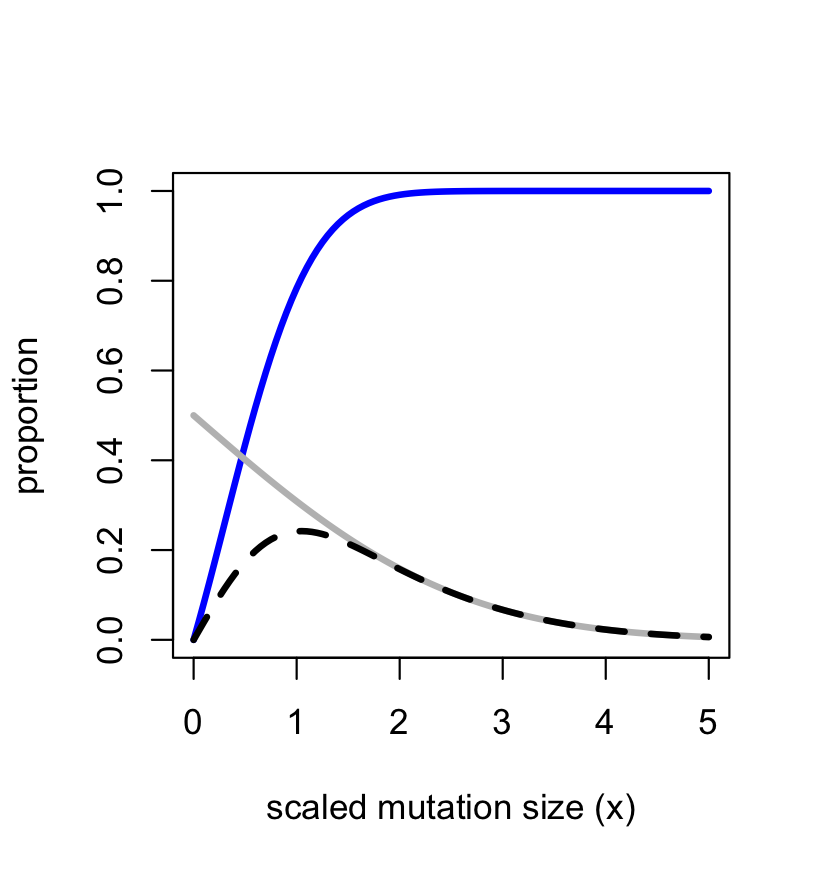

The results clearly show that beneficial mutations with small, scaled sizes (*i.e.*, *x* << 1) are not expected to meet conditions for balancing selection. Rather, they experience positive selection and are favoured to sweep to fixation. In contrast, beneficial mutations of moderate-to-large size (*x* > 1) are likely to experience balancing selection, at least in the short term.

Whether or not balancing selection is prevalent will depend on the distribution of mutation sizes (the distribution of *x*). If the distribution of *x* is concentrated near zero—which we can think of as the infinitesimal limit—positive selection prevails. If, however, the distribution of *x* is broad and spans small through large mutations, then (short-term) balancing selection should be common. For purpose of illustration, if we assume that *x* is approximately uniform across the span of *x* where beneficial mutations are likely (*i.e.*, 0 < *x* < 5), then the total proportion of beneficial mutations that exhibits short-term balancing selection will be:

$$\begin{aligned} f_{bal}\approx\frac{\int_{0}^{\infty} \Pr\left( \text{bal.}|x \right)dx}{\int_{0}^{\infty} \Pr\left( \text{ben.}|x \right)dx}=\frac{2}{3} .\#\left( 54 \right) \end{aligned}$$

Of course, only a fraction of beneficial mutations will contribute to adaptation, since many may be lost due to drift, particularly if their beneficial effects are small. If we exclusively focus on mutations that successfully invade the population, then our quantitative predictions about balancing selection somewhat change (see McDonough and Connallon 2023; McDonough et al. 2024). The probability that a random mutation of size *x* is both beneficial and successfully invades the population (*i.e.*, its probability of establishment) is:

$$\begin{aligned} \Pr\left( \text{est.}|x \right)=\frac{z^{2}}{n}x\left( \sqrt{\frac{2}{\pi}}\exp\left( -\frac{1}{8}x^{2} \right)-\frac{1}{2}x\left( 1-\mathrm{erf} \left( \frac{x}{2\sqrt{2}} \right) \right) \right) .\#\left( 55 \right) \end{aligned}$$

The probability of establishment for mutations subject to balancing selection is:

$$\Pr\left( \text{est. bal.}|x \right)=\frac{z^{2}}{n}x\sqrt{\frac{2}{\pi}}\left( \exp\left( -\frac{1}{8}x^{2} \right)-\exp\left( -\frac{9}{8}x^{2} \right) \right)$$

$$\begin{aligned} -\frac{z^{2}}{4n}x^{2}\left( \mathrm{erf} \left( \frac{3x}{2\sqrt{2}} \right)-\mathrm{erf} \left( \frac{x}{2\sqrt{2}} \right) \right) .\#\left( 56 \right) \end{aligned}$$

The following figure compares the two establishment probabilities ($\Pr\left( \text{est.}|x \right)$ in grey; $\Pr\left( \text{est. bal.}|x \right)$ in black; each is divided by *z*^2^/*n* for ease of comparison) and their ratio (blue).

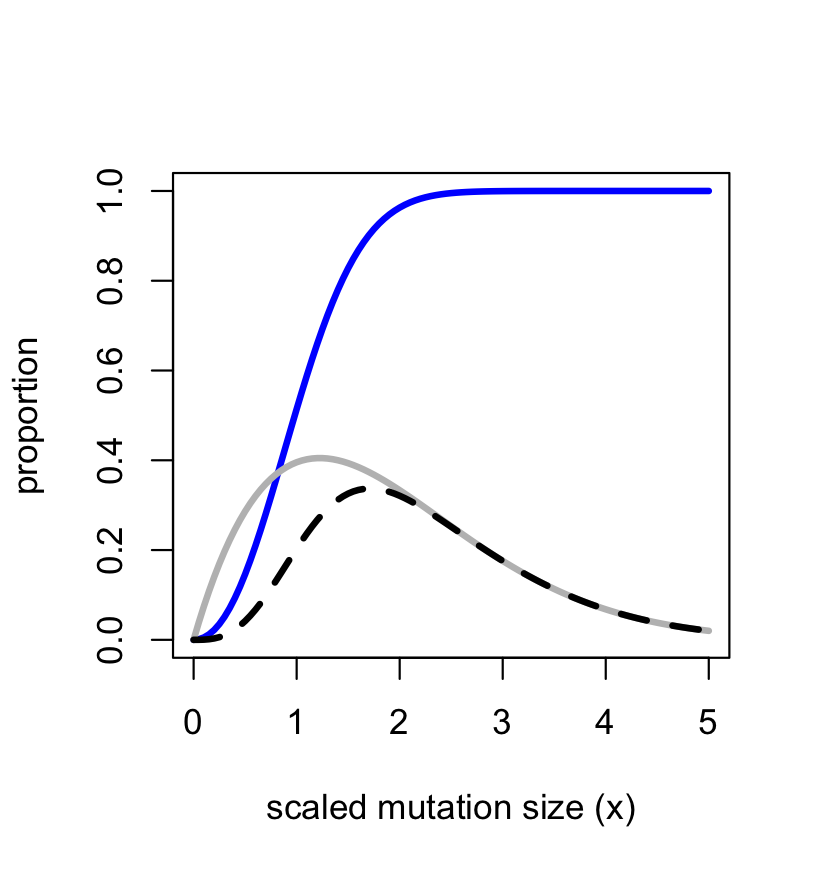

Compared to new mutations, the curves of established mutations are shifted to the right, which reflects that mutations with small phenotypic effects are unlikely to become established because they are, at most, weakly beneficial and often lost due to drift (see Orr 1998). For an invading mutation whose effect size is small, the probability of balancing selection is small, whereas large effect invading mutations should often be subject to balancing selection. If we assume a broad distribution of mutant effect sizes (again, *x*~uniform across the interval where invasion is probable), then the proportion of invading mutations that are subject to short-term balancing selection becomes:

$$\begin{aligned} f_{bal}\approx\frac{\int_{0}^{\infty} \Pr\left( \text{bal.}|x \right)dx}{\int_{0}^{\infty} \Pr\left( \text{ben.}|x \right)dx}=\frac{20}{27}\approx0.74 .\#\left( 57 \right) \end{aligned}$$

***Adaptive walks.*** Simulations of multi-step adaptive walks to an optimum, with each step involving the invasion of a new mutation into the population, predict that outcrossing diploid populations adapt through a series of partial selective sweeps, many of which result in a new balanced polymorphic state that persists until the next mutation invades and perturbs the polymorphic system (Sellis et al. 2011). Because the distance of the population to the optimum progressively shrinks over time (*z* declines during adaptation), whereas trait dimensionality and the distribution of raw mutation sizes do not change (*n* and the distribution of *r* is constant), mutation sizes in Fisher’s scale will increase during the adaptive walk, which makes balancing selection increasingly likely as long as the population remains displaced from its optimum. While balanced states are common during the adaptive walk, no single state is indefinitely maintained. In other words, long-term balancing selection is not predicted by this model.

**Models of stabilizing selection on polygenic traits**

Balancing selection is a transient phenomenon in the Fisher’s geometric model framework, and each polymorphic state persists until a new beneficial mutation invades the population and displaces it. This framework is, therefore, best suited to modelling evolution of traits that are somewhat mutation-limited and evolve in the manner of stepwise adaptive walks.

For some traits—particularly those with a large mutational target in which variation is highly polygenic (e.g., body size is a well-studied example)—this framework is clearly unlikely to apply. Here, standing genetic variation should be sufficient for the trait to quickly evolve toward the optimum, and opportunities for even transient balanced polymorphic states should be limited. Consider, for example, the basic framework set out in Turelli and Barton (2004), which describes the evolutionary dynamics of a single quantitative trait whose expression is affected by *L* bi-allelic loci. The set of loci are unlinked and segregate independently of one another. Effects of the alleles on the trait are additive both within and among loci. We will assume the population is large (we will ignore drift), diploid, and randomly mating.

At the *i*^th^ locus, allele *A_i_* has a population frequency of *p_i_*. Each copy of the allele causes an individual’s trait expression to increase by amount $\alpha_{i}$ ($\alpha_{i}>0$). The alternate allele, *a_i_*, is at frequency *q_i_* and decreases trait expression by the same magnitude (see Table S2).

| **Table S2. Effect of the *i*^th^ locus on the expression of a continuous trait** | | | |
| --- | --- | --- | --- |
|  |  | ***Genotype*** |  |
|  | ***A_i_A_i_*** | ***A_i_a_i_*** | ***a_i_a_i_*** |
| **Phenotypic effect:** | $\alpha_{i}$ | 0 | $-\alpha_{i}$ |
| **Zygote frequency:** | $p_{i}^{2}$ | $2p_{i}q_{i}$ | $q_{i}^{2}$ |

We will assume for simplicity that an individual’s trait expression value depends on its breeding value (multi-locus genotype) and that environmental effects on trait expression are negligible. The relative fitness of an individual with breeding value *G* follows the Gaussian function:

$$\begin{aligned} w\left( G \right)=\exp\left( -\frac{\left( G-\theta\right)^{2}}{2V_{S}} \right) ,\#\left( 58 \right) \end{aligned}$$

where $\theta$ is the optimal expression value, and $V_{S}$ is the width of the fitness function. If the number of segregating loci (*L*) is large and the phenotypic effects of individual genetic variants are small (variation is sufficiently fine-grained), then *G* should be normally distributed among members of the population, and mean relative fitness among will be:

$$\begin{aligned} \bar{w}=\int w\left( G \right)f\left( G \right)dG=\frac{\sqrt{V_{S}}}{\sqrt{V_{S}+V_{A}}}\exp\left( -\frac{\left( \theta-\mu\right)^{2}}{2V_{S}+2V_{A}} \right) ,\#\left( 59 \right) \end{aligned}$$

where $\mu=\sum_{i=1}^{L} \alpha_{i}\left( p_{i}-q_{i} \right)$ is the trait mean for the population and $V_{A}=2\sum_{i=1}^{L} \alpha_{i}^{2}p_{i}q_{i}$ is the additive genetic variance for the trait.

Assuming no linkage disequilibria among loci, the evolutionary dynamics of the *i*^th^ polymorphic locus are described by:

$$\begin{aligned} \Delta p_{i}=\frac{p_{i}\left( 1-p_{i} \right)}{2}\frac{\partial\log\left( \bar{w} \right)}{\partial p_{i}}=\frac{p_{i}\left( 1-p_{i} \right)}{2\left( V_{S}+V_{A} \right)}\left[ \left( \theta-\mu\right)\frac{\partial\mu}{\partial p_{i}}+\frac{1}{2}\left( \frac{\left( \theta-\mu\right)^{2}}{V_{S}+V_{A}}-1 \right)\frac{\partial V_{A}}{\partial p_{i}} \right] ,\#\left( 60 \right) \end{aligned}$$

where $\frac{\partial\mu}{\partial p_{i}}=2\alpha_{i}$ and $\frac{\partial V_{A}}{\partial p_{i}}=2\alpha_{i}^{2}\left( 1-2p_{i} \right)$. Substituting the latter two identities gives us:

$$\begin{aligned} \Delta p_{i}=\frac{p_{i}\left( 1-p_{i} \right)}{2\left( V_{S}+V_{A} \right)^{2}}\left[ 2\alpha_{i}\left( \theta-\mu\right)\left( V_{S}+V_{A} \right)+\alpha_{i}^{2}\left( 1-2p_{i} \right)\left( \left( \theta-\mu\right)^{2}-V_{S}-V_{A} \right) \right] .\#\left( 61 \right) \end{aligned}$$

This result shows that selection at each locus should have both a directional and a frequency-dependent component, as captured (respectively) by the first and second terms in the square brackets. The first term (the directional selection component: $2\alpha_{i}\left( \theta-\mu\right)\left( V_{S}+V_{A} \right)$) promotes the spread of alleles that shift the mean phenotype towards the optimum. In our case, since $\alpha_{i}>0$, the directional selection component favours the *A_i_* allele when the population mean is below optimum ($\theta>\mu$), and it favours the *a_i_* allele when the population is above optimum ($\theta<\mu$). The second term in square brackets is frequency-dependent and arises from stabilizing selection to reduce the trait variance. In particular, the second term favours elimination of the rarer of the two alleles (the minor allele), which reduces *V_A_*.

When the frequency-dependent term dominates, selection is variance reducing, which prohibits balancing selection at the locus. For example, in a population whose trait mean has evolved to match the optimum ($\mu=\theta$), the predicted change in allele frequency simplifies to:

$$\begin{aligned} \Delta p_{i}=\frac{\alpha_{i}^{2}p_{i}\left( 1-p_{i} \right)\left( 2p_{i}-1 \right)}{2\left( V_{S}+V_{A} \right)}\#\left( 62 \right) \end{aligned}$$

and selection therefore favours a decline in the *A_i_* allele frequency when *p_i_* < 0.5, and it favours an increase in frequency when *p_i_* > 0.5. However, either change would cause a displacement of the population’s trait mean from the optimum, which would generate some degree of directional selection on the trait and its underlying polymorphic loci. In the long run, if there is a homozygous multi-locus genotype that yields a close match between the trait mean and the optimum, then at equilibrium, all *L* loci will experience purifying selection (against the minor allele), and balancing selection will not occur. If, however, the best homozygous genotype remains displaced from the optimum, then a locus with a sufficiently large phenotypic effect can potentially remain polymorphic owing to balancing selection, and such a polymorphism can be maintained indefinitely as long as the mutational architecture of the trait remains static (see Turelli and Barton 2004; Flintham et al. 2024).

**Genetic variation maintained by pleiotropic balancing selection**

A variety of authors have considered the possibility that balanced polymorphisms might affect variation in a quantitative trait that is subject to stabilizing selection, but balancing selection arises from the effect of the locus on another component of fitness. In these models, stabilizing selection on the quantitative trait tends to favour the removal of the polymorphism, while balancing selection through the other fitness component favours the maintenance of the polymorphism. The overall effect of the two episodes of selection is that the polymorphism may be maintained if balancing selection is stronger than the effect of stabilizing selection, or it may be lost if stabilizing selection overpowers balancing selection. This scenario has been modelled, in different forms, by Robertson (1956), Bulmer (1973), Gillespie (1984), Barton (1990) and Turelli and Barton (2004). Our derivation (below) differs in some ways from theirs, but the result captures the essence of these models.

Consider a model in which there is viability selection with respect to the basic molecular function of a given gene, and then there is a subsequent episode of selection on the quantitative trait whose expression is affected by the focal gene and various other polymorphic genes. Since the mutational target of a single gene is small, variation affecting its molecular function is decidedly non-continuous and is reasonably likely to be dominated by a small number of variants. For example, it may be segregating for one favourable variant at a given time, or subject to balancing selection in a manner described by classical population genetic models of balancing selection.

Suppose there is a pair of variants at our focal locus: allele *A*_1_ is at frequency *p* and allele *A*_2_ is at frequency *q* = 1 – *p*. The relative effects of the three genotypes on the first episode of selection are *w*_11_ for *A*_1_*A*_1_, *w*_12_ for *A*_1_*A*_2_, and *w*_22_ for *A*_2_*A*_2_. Assuming that offspring are produced through random mating of the parents of the previous generation, then the frequencies of the three genotypes after the first round of selection are:

$$\begin{aligned} f_{11}=\frac{p^{2}v_{11}}{p^{2}v_{11}+2pqv_{12}+q^{2}v_{22}} ,\#\left( 63a \right) \end{aligned}$$

$$\begin{aligned} f_{12}=\frac{2pqv_{12}}{p^{2}v_{11}+2pqv_{12}+q^{2}v_{22}} ,\#\left( 63b \right) \end{aligned}$$

$$\begin{aligned} f_{22}=\frac{q^{2}v_{22}}{p^{2}v_{11}+2pqv_{12}+q^{2}v_{22}} .\#\left( 63c \right) \end{aligned}$$

The quantitative trait is under stabilizing selection to an optimum. Each polymorphic locus contributes additively to the trait’s expression. The phenotypic effects of the focal locus are:

|  | ***A*_1_*A*_1_** | ***A*_1_*A*_2_** | ***A*_2_*A*_2_** |
| --- | --- | --- | --- |
| **Avg. phenotypic effect** | $-\alpha$ | 0 | $\alpha$ |
| **Frequency** | *f*_11_ | *f*_12_ | *f*_22_ |

The trait variance due to other polymorphic loci and random environmental factors is given by $V_{R}+V_{E}$, where $V_{R}$ is the residual genetic variance due to the other loci, and *V_E_* is the environmental variance. The total phenotypic variance of the trait (including the focal locus) will be $V_{P}=V_{A}+V_{R}+V_{E}$, where $V_{A}$ is the additive genetic variance attributable to the focal locus. The phenotypes associated with each genotype for the focal locus are each drawn from a normal distribution. The phenotype of an individual with focal genotype *ij* is drawn from a normal distribution with genotype-specific mean and variance $V_{R}+V_{E}$. With $\mu$ representing the mean breeding value with respect to all loci except the focal locus, the mean for individuals with genotype *A*_1_*A*_1_, *A*_1_*A*_2_, and *A*_2_*A*_2_ will be (respectively) $\mu-\alpha$, $\mu$, and $\mu+\alpha$.

Fitness with respect to the second episode of selection is a Gaussian function of trait expression (*z*):

$$\begin{aligned} w\left( z \right)=\exp\left( -\frac{z^{2}}{2V_{S}} \right) .\#\left( 64 \right) \end{aligned}$$

Therefore, the average fitness associated with individuals with each of the three focal genotypes is (*A*_1_*A*_1_, *A*_1_*A*_2_, and *A*_2_*A*_2_) will be:

$$\begin{aligned} W_{11}=\frac{1}{\sqrt{2\pi\left( V_{R}+V_{E} \right)}}\int_{-\infty}^{\infty} \exp\left( -\frac{\left( z-\mu+\alpha\right)^{2}}{2V_{R}+2V_{E}}-\frac{z^{2}}{2V_{S}} \right)dz\#\left( 65a \right) \end{aligned}$$

$$=\frac{\sqrt{V_{S}}}{\sqrt{V_{R}+V_{E}+V_{S}}}\exp\left( -\frac{\left( \mu-\alpha\right)^{2}}{2V_{R}+2V_{E}+2V_{S}} \right) ,$$

$$\begin{aligned} W_{12}=\frac{1}{\sqrt{2\pi\left( V_{R}+V_{E} \right)}}\int_{-\infty}^{\infty} \exp\left( -\frac{\left( z-\mu\right)^{2}}{2V_{R}+2V_{E}}-\frac{z^{2}}{2V_{S}} \right)dz\#\left( 65b \right) \end{aligned}$$

$$=\frac{\sqrt{V_{S}}}{\sqrt{V_{R}+V_{E}+V_{S}}}\exp\left( -\frac{\mu^{2}}{2V_{R}+2V_{E}+2V_{S}} \right) ,$$

$$\begin{aligned} W_{22}=\frac{1}{\sqrt{2\pi\left( V_{R}+V_{E} \right)}}\int_{-\infty}^{\infty} \exp\left( -\frac{\left( z-\mu-\alpha\right)^{2}}{2V_{R}+2V_{E}}-\frac{z^{2}}{2V_{S}} \right)dz\#\left( 65c \right) \end{aligned}$$

$$=\frac{\sqrt{V_{S}}}{\sqrt{V_{R}+V_{E}+V_{S}}}\exp\left( -\frac{\left( \mu+\alpha\right)^{2}}{2V_{R}+2V_{E}+2V_{S}} \right) .$$

The frequency of the *A*_2_ allele after selection through the second fitness component is:

$$\begin{aligned} q^{'}=\frac{f_{22}W_{22}+\frac{1}{2}f_{12}W_{12}}{f_{22}W_{22}+f_{12}W_{12}+f_{11}W_{11}}=\frac{q^{2}v_{22}W_{22}+pqv_{12}W_{12}}{q^{2}v_{22}W_{22}+2pqv_{12}W_{12}+p^{2}v_{11}W_{11}} .\#\left( 66 \right) \end{aligned}$$

This completes the life cycle of a single generation, so that *A*_1_ is expected to increase over time when $q^{'}>q$, it is expected to decrease over time when $q^{'}<q$, and the system is at an equilibrium when $q^{'}=q$.

The trait mean before the second episode of selection is:

$$\begin{aligned} \bar{z}=\mu+\alpha\left( f_{22}-f_{11} \right)=\mu+\alpha\left( \frac{q^{2}v_{22}-p^{2}v_{11}}{p^{2}v_{11}+2pqv_{12}+q^{2}v_{22}} \right) .\#\left( 67 \right) \end{aligned}$$

Let’s assume that the residual mean breeding value ($\mu$) can shift rapidly relative to the rate of change of the alleles at the focal locus. In this case, we can justify a separation of timescales for the evolution of $\mu$ (fast) and *p* and *q* (slow), and the mean breeding value for background loci ($\mu$) should therefore evolve to a *quasi*-equilibrium state that is a function of the allele frequencies of the focal locus. The quasi-equilibrium breeding value $\tilde{\mu}=\alpha\left( f_{11}-f_{22} \right)$ causes the trait mean to match the trait optimum ($\bar{z}=0$ in this case). The quasi-equilibrium fitness associated with the focal genotypes for the second episode of selection will be:

$$\begin{aligned} \tilde{W}_{11}=\frac{\sqrt{V_{S}}}{\sqrt{V_{R}+V_{E}+V_{S}}}\exp\left( -\frac{{\alpha^{2}\left( f_{11}-f_{22}-1 \right)}^{2}}{2V_{R}+2V_{E}+2V_{S}} \right),\#\left( 68a \right) \end{aligned}$$

$$\begin{aligned} \tilde{W}_{12}=\frac{\sqrt{V_{S}}}{\sqrt{V_{R}+V_{E}+V_{S}}}\exp\left( -\frac{{\alpha^{2}\left( f_{11}-f_{22} \right)}^{2}}{2V_{R}+2V_{E}+2V_{S}} \right),\#\left( 68b \right) \end{aligned}$$

$$\begin{aligned} \tilde{W}_{22}=\frac{\sqrt{V_{S}}}{\sqrt{V_{R}+V_{E}+V_{S}}}\exp\left( -\frac{{\alpha^{2}\left( f_{11}-f_{22}+1 \right)}^{2}}{2V_{R}+2V_{E}+2V_{S}} \right). \#\left( 68c \right) \end{aligned}$$

The condition for a global protected polymorphism at the focal locus occurs when both boundary equilibria are unstable. Evaluating at the quasi-equilibrium for each boundary equilibrium of the focal locus (*q* = 0 and *q* = 1, respectively), we have:

$$\begin{aligned} \left. \frac{dq^{'}}{dq} \right|_{q=0}=\frac{v_{12}\tilde{W}_{12}}{v_{11}\tilde{W}_{11}}=\frac{v_{12}}{v_{11}}\exp\left( -\frac{\alpha^{2}}{2V_{R}+2V_{E}+2V_{S}} \right),\#\left( 69a \right) \end{aligned}$$

$$\begin{aligned} \left. \frac{dq^{'}}{dq} \right|_{q=1}=\frac{v_{12}\tilde{W}_{12}}{v_{22}\tilde{W}_{22}}=\frac{v_{12}}{v_{22}}\exp\left( -\frac{\alpha^{2}}{2V_{R}+2V_{E}+2V_{S}} \right),\#\left( 69b \right) \end{aligned}$$

and the focal locus is subject to balancing selection (there is a global protected polymorphism) when the conditions $\left. \frac{dq^{'}}{dq} \right|_{q=0}>1$ and $\left. \frac{dq^{'}}{dq} \right|_{q=1}>1$ are both true. In other words, the polymorphism will be globally protected under the condition:

$$\begin{aligned} \min\left( \frac{v_{12}}{v_{11}},\frac{v_{12}}{v_{22}} \right)>\exp\left( \frac{\alpha^{2}}{2V_{R}+2V_{E}+2V_{S}} \right).\#\left( 70 \right) \end{aligned}$$

For example, in the case of heterozygote advantage through the first fitness component and unit variance for the trait $V_{R}+V_{E}=1$, we have:

$$\begin{aligned} \min\left( s_{1},s_{2} \right)>1-\exp\left( -\frac{\alpha^{2}}{2\left( 1+V_{S} \right)} \right),\#\left( 71 \right) \end{aligned}$$

where $v_{11}=1-s_{1}$, $v_{12}=1$, and $v_{22}=1-s_{2}$ (which conforms to the heterozygote advantage model from Box 1 in the main text).

In contrast to the classic model of heterozygote advantage at a single locus, where all positive values of $s_{1}$ and $s_{2}$ ($0<s_{1},s_{2}\leq1$) will generate balancing selection in a randomly mating population, stabilizing selection on the quantitative trait potentially overpowers balancing selection through the first fitness component. Thus, whereas 100% of the parameter space of heterozygote advantage can generate balancing selection in the classical model, the proportion of parameter space that leads to a global protected polymorphism in the pleiotropic stabilizing selection model is:

$$\begin{aligned} \exp\left( -\frac{\alpha^{2}}{1+V_{S}} \right).\#\left( 72 \right) \end{aligned}$$

For the special case in which there is sufficiently strong and symmetric heterozygote advantage through the first fitness component (i.e., $s_{1}=s_{2}>1-\exp\left( -\frac{\alpha^{2}}{2\left( 1+V_{S} \right)} \right)$), the equilibrium allele frequencies at the focal locus will be intermediate ($\hat{p}=\hat{q}=0.5$), and the contribution of the locus to the trait’s genetic variance will be:

$$\begin{aligned} V_{A}=2\alpha^{2}\hat{p}\hat{q}=\frac{\alpha^{2}}{2}.\#\left( 73 \right) \end{aligned}$$

At equilibrium, the total genetic variance for the trait would be $V_{G}=V_{A}+V_{R}$, and it is certainly possible that the balanced polymorphism might account for a large fraction of the overall variance of the trait even though selection on that trait does not explain the maintenance of the polymorphism because balancing selection arises from selection through the other fitness component.

**Summary of balancing selection and genetic variation affecting continuous traits**

In the absence of pleiotropy between different components of fitness, conditions for long-term balancing selection can become restrictive, particularly as the trait mean of the population approaches the trait optimum. In contexts where adaptation is mutation-limited—which may depend on context of the traits or the populations that are responding to selection (Rousselle et al. 2020)—we may see a series of transient balanced polymorphic states at loci contributing to variation in traits selected to an optimum. If there are homozygous genotypes that allow the population to reach the optimum, their fixation limits opportunities for long-term stabilizing selection, since at optimum stabilizing selection becomes variance reducing and disfavours rare variants.

In cases where variants affecting a polygenic quantitative trait have pleiotropic effects on other fitness components, balancing selection through the other component can maintain polymorphism affecting the trait even if selection on the trait itself is not conducive to balancing selection. This scenario seems particularly likely when for loci whose pleiotropic effects occur in traits with a relatively simple mutational architecture (e.g., simple molecular functions in which there is no continuum of allelic effects on the trait in question).
