## Appendix 5 (Citation analysis) for "A century of theories of balancing selection"

**Appendix 5. Publication patterns in balancing selection research**

**Data collection and curation**

**Web of Science bibliography**

Empirical papers on balancing selection were compiled from the Web of Science Core Collection database (<https://www.webofscience.com/wos/woscc>) on 11/09/2023, using search terms "balancing selection", "balanced polymorphism", and "protected polymorphism", resulting in 2936 records. The term "stable polymorphism" was added on 02/11/2023, resulting in 109 additional records.

Records were reviewed and curated using Rayyan (https://rayyan.ai). Each paper was evaluated by one of four reviewers, and most papers were reviewed by a single reviewer. Papers were only included if they identified or discussed balancing selection at a specific locus (or loci). We removed duplicates, and excluded papers focusing on other forms of selection, or otherwise tangential (e.g., discussing “immunity loci” without characterizing forms of selection). Each paper was categorized as a review, theoretical, or empirical study. A total of 996 papers were thus obtained, of which 872 were empirical, 60 were reviews and the remaining were theory. Theory studies formed a small minority of papers found in this manner. We therefore used a separate process to compile the theoretical bibliography (see ‘Manually curated bibliography of theory papers’). For each paper, where possible, we noted information on the scenario of balancing selection, which was labelled as follows (brackets indicate terminology used in the main text): HA (heterozygote advantage), FD (negative frequency-dependent selection), MD (meiotic drive), AP (antagonistic pleiotropy), SA (sexually antagonistic selection), TF (temporally fluctuating selection), NA (niche antagonism), PA (ploidally antagonistic selection)).

**Manually curated bibliography of theory**

To complement the Web of Science bibliography, which predominantly identified empirical and review papers, we also performed a manual search for theory papers on balancing selection. The procedure was to go through bibliographies of known balancing theory papers and add papers that modelled conditions for balancing selection. For logistical reasons, book chapters were excluded (access rights were often difficult to come by), as were papers on multi-allelic balancing selection (not a major focus of our review). This search resulted in 406 papers. We noted the scenario of balancing selection (categories as above), and the aspect of biological complexity (e.g., if a paper modelled haploid balancing selection in a finite population, it was categorized as ‘finite population’ and ‘haploids and haplo-diploids’).

**Data processing and data availability**

We used RStudio (RStudio Team, 2020) to manipulate and visualise data, using packages dlpyr (Wickham *et al.*, 2023), tidyverse (Wickham *et al.*, 2019), tidygraph (Pedersen, 2024), ggplot2 (Wickham, 2016), openalexR (Aria *et al.*, 2024), svglite (Wickham *et al.*, 2024). Full documentation and code underlying figures can be found at: <https://github.com/marzw/BalancingSelection_STN/>

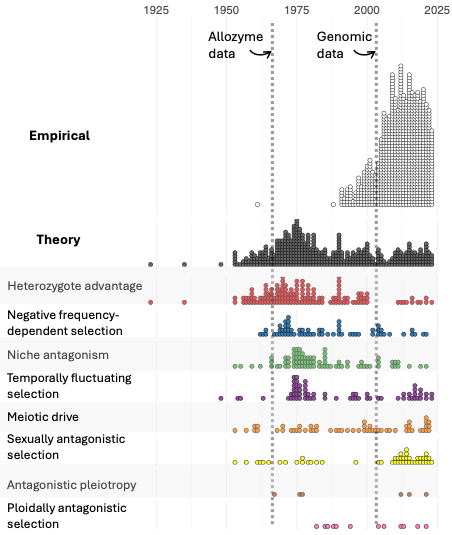

A timeline of balancing selection theory and data. Each dot represents a paper on balancing selection (N = 872 empirical papers, N = 402 theory papers). The top two panels are identical to those presented in Figure 1 of the main text. The bottom panels zoom in on the theoretical papers and separate them into different scenarios of balancing selection.

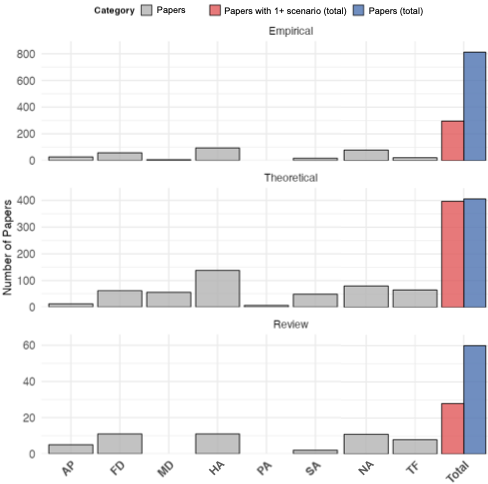

Number of publications for each different scenario of balancing selection (grey), and across scenarios (red, blue), for each category of paper (empirical, theoretical, review). AP = antagonistic pleiotropy, FD = negative frequency-dependent selection, MD = meiotic drive, HA = heterozygote advantage, PA = ploidally antagonistic selection, SA = sexually antagonistic selection, NA = niche antagonism, TF = temporally fluctuating selection.

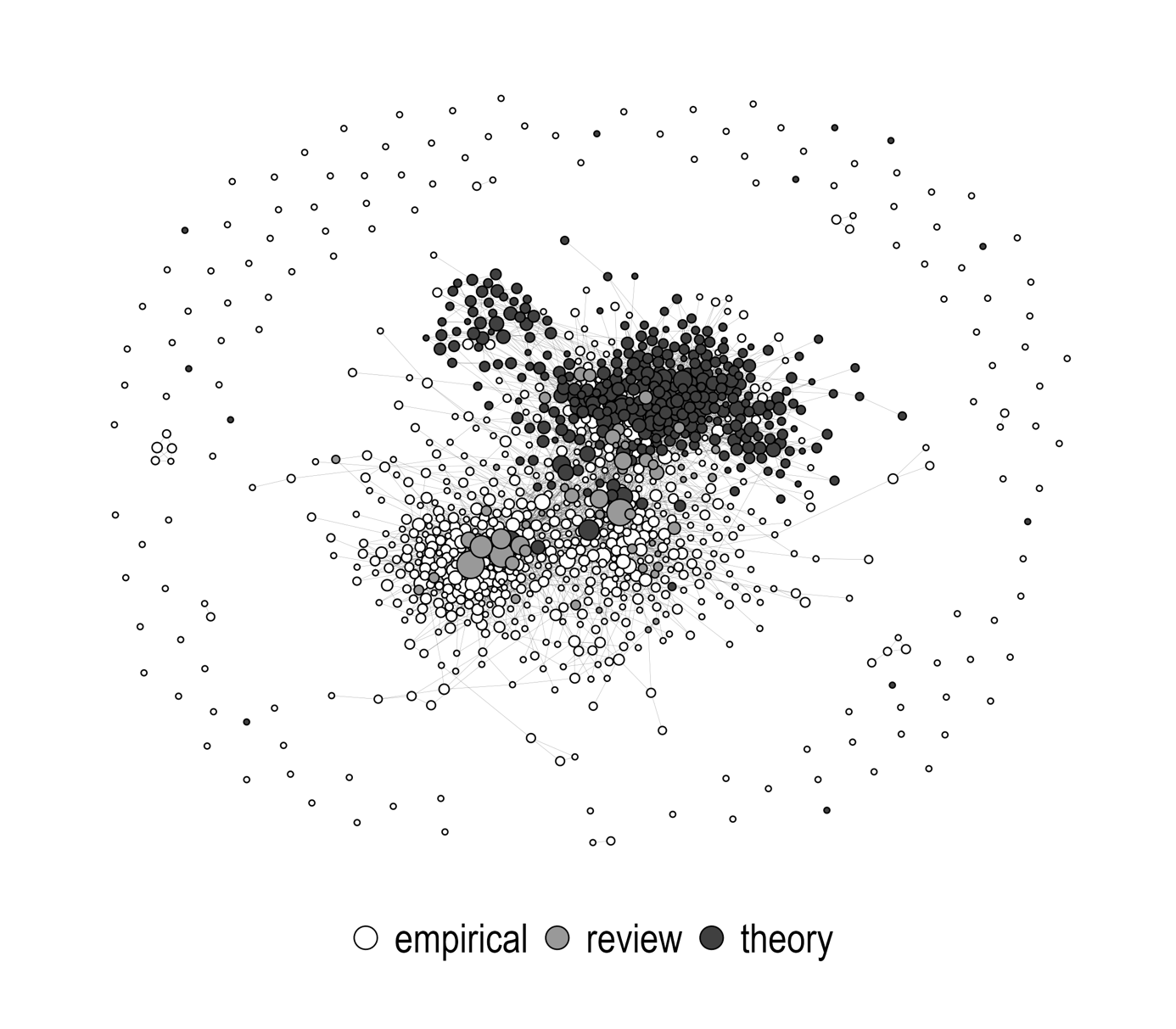

Local citation network of research on balancing selection. Each node (circle) represents one article in the dataset (N = 1107). Node colour denotes article type (empirical, theory, review). Node size is proportional to number of times the article is cited locally within the network (not global citation count). Edges show citations within the local network (N = 4358). Unconnected nodes do not cite articles within the network and are not cited by articles within the network. Only articles with a DOI are included.

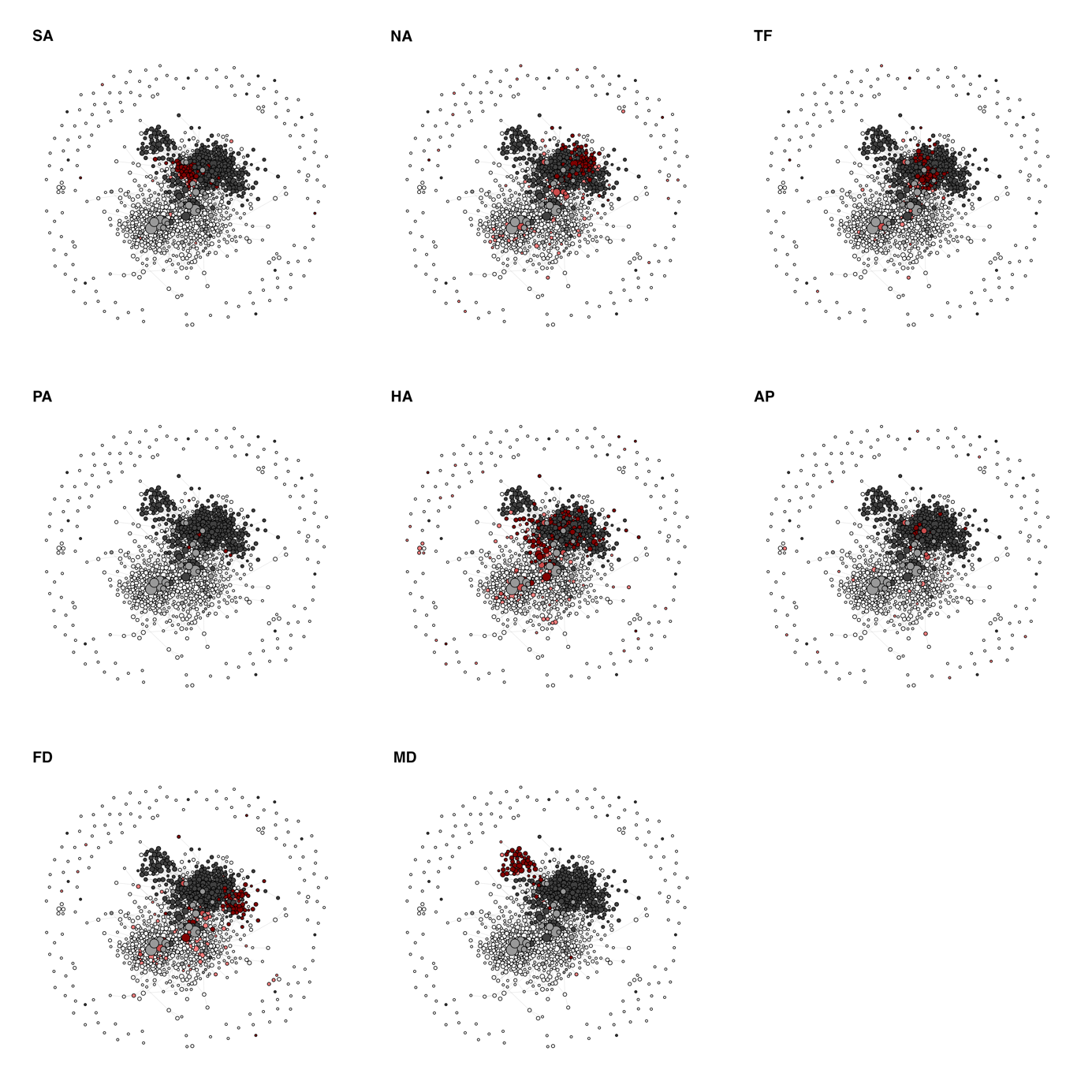

Local citation network of research on balancing selection. Each node (circle) represents one article in the dataset (N = 1107). Node shade denotes article type (empirical, theory, review), and red colouration highlights the indicated scenario (SA = sexually antagonistic selection, NA = niche antagonism; TF = temporally fluctuating selection, PA = ploidally antagonistic selection, HA = heterozygote advantage, AP = antagonistic pleiotropy, FD = negative frequency-dependent selection, MD = meiotic drive).

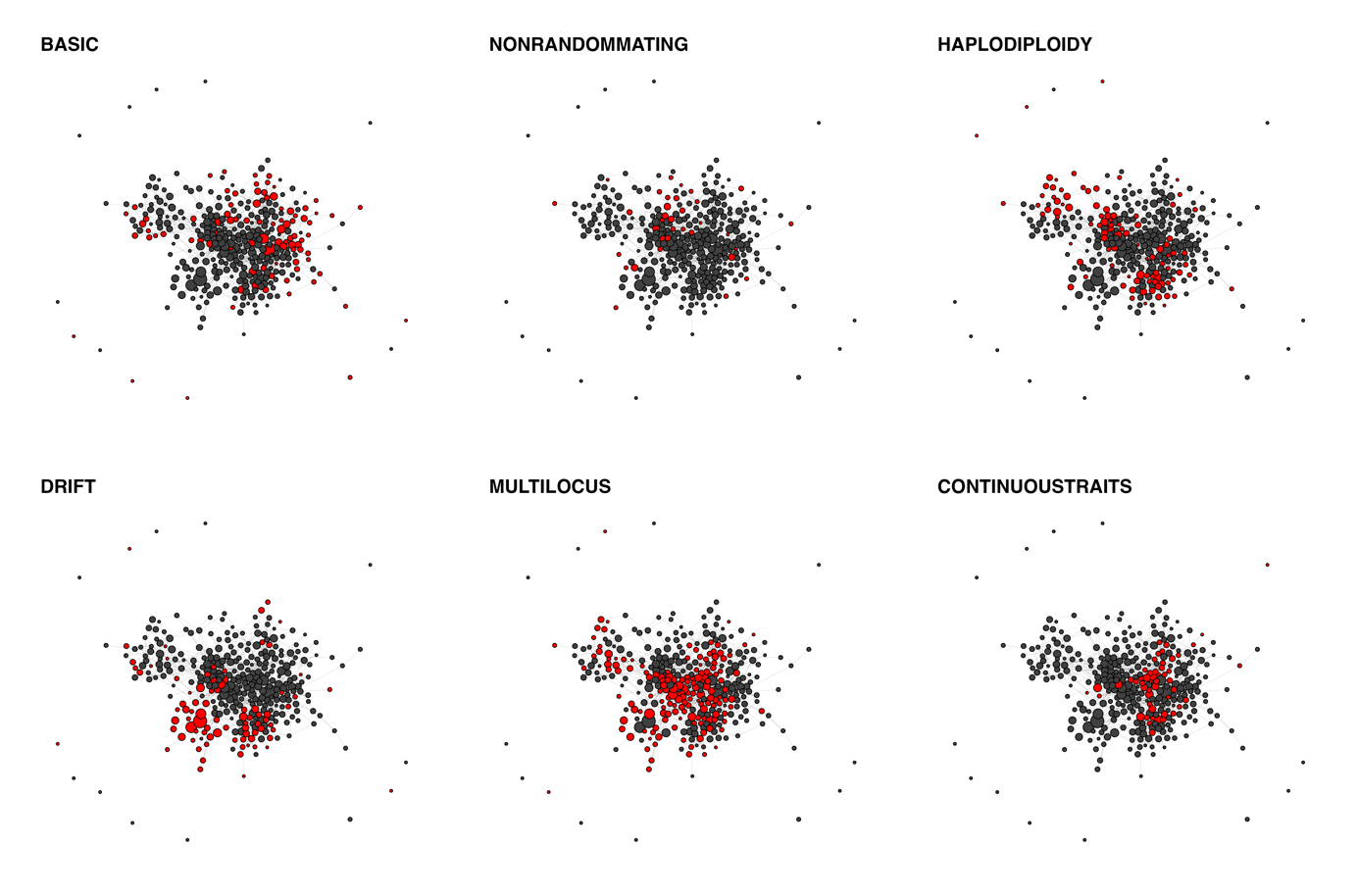
 Local citation network of theoretical research on balancing selection. Each node (circle) represents one article in the dataset (N = 334). Red colour highlights the indicated model feature. BASIC (the idealized model we present in ‘Models of balancing selection: the basics’), NONRANDOMMATING (‘Non-random mating’), HAPLODIPLOIDY (‘Haploids and haplo-diploids’), DRIFT (‘Finite populations’), MULTILOCUS (‘Multi-locus systems with linkage’), CONTINUOUSTRAITS (‘Traits selected towards an optimum’).

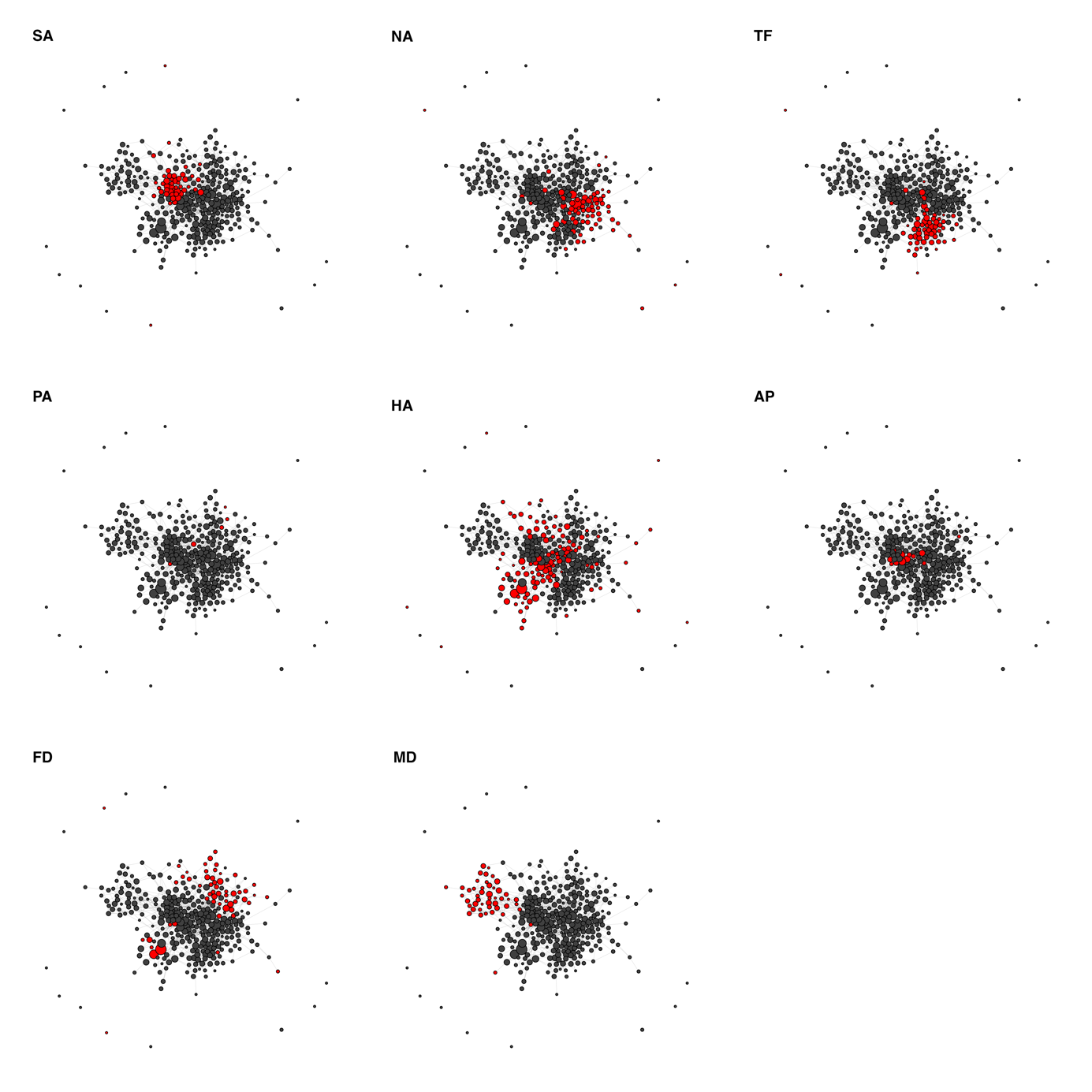

Local citation network of theoretical research on balancing selection. Each node (circle) represents one article in the dataset (N = 334). Red colour highlights the indicated scenario. (SA = sexually antagonistic selection, NA = niche antagonism; TF = temporally fluctuating selection, PA = ploidally antagonistic selection, HA = heterozygote advantage, AP = antagonistic pleiotropy, FD = negative frequency-dependent selection, MD = meiotic drive).

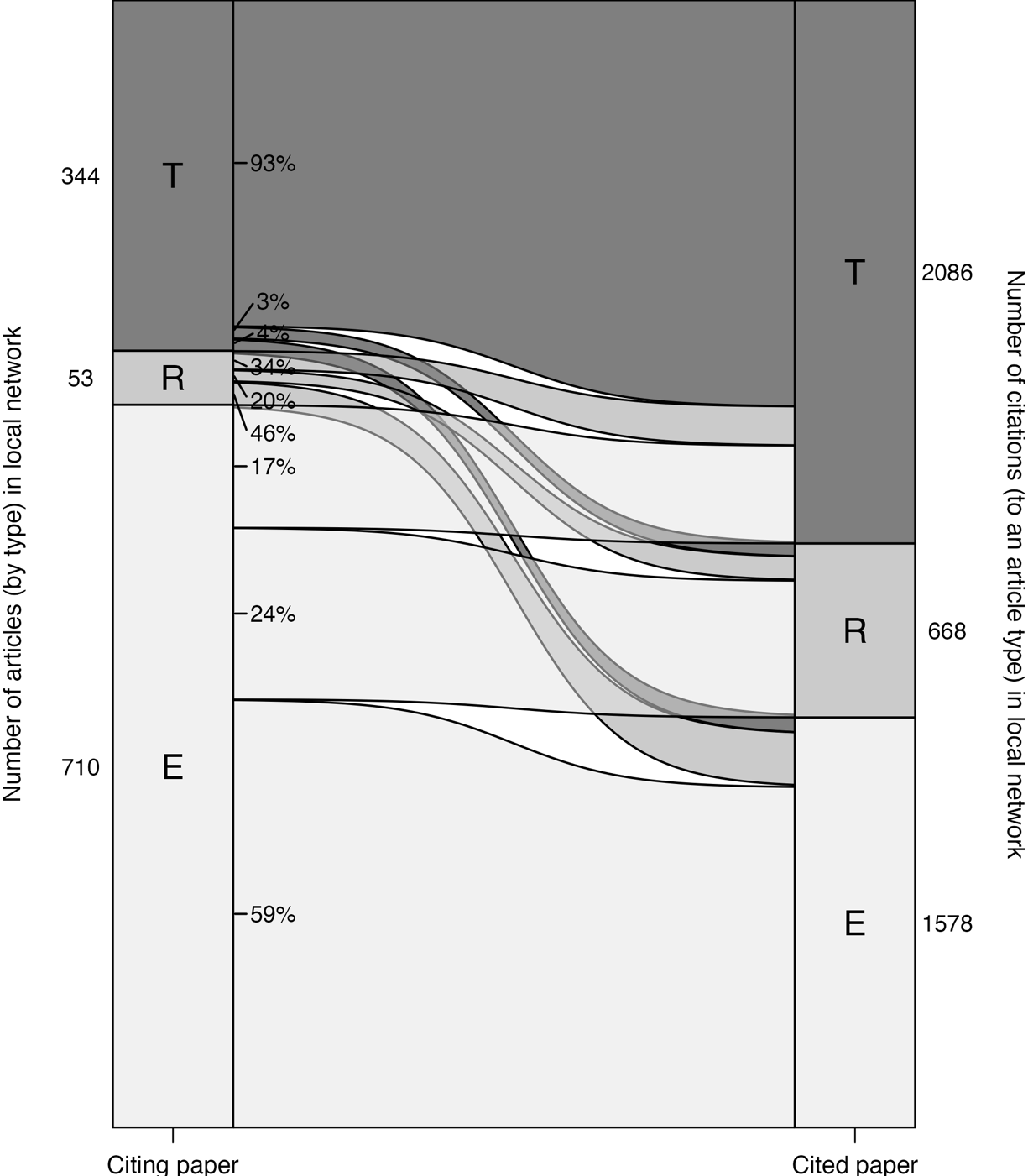

Citation patterns for each article type (E = empirical, R = review, T = theoretical) within the local network. The boxes on the left (“Citing paper”) are scaled proportional to the number of articles of a given type in the network (shown in the adjacent number). The boxes on the right (“Cited paper”) are scaled proportional to the number of times an article of a given type is cited in the network (shown in the adjacent number). The flows between the two sides have heights proportional to the proportion of citations from a given article type to another article type. For example, the topmost flow, from T to T, shows that 93% of citations by theory papers are to other theory papers, whereas citations to review papers (T to R) and empirical papers (T to E) make up only 3% and 4%, respectively.

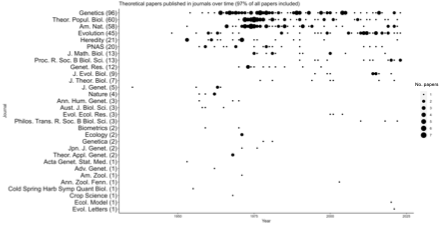

Timeline of publication of theoretical papers on balancing selection, split by journal.

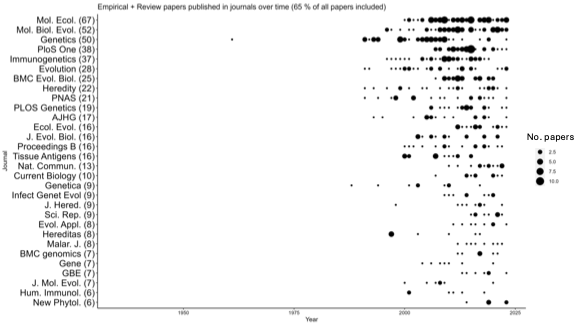

Timeline of publication of empirical and review papers on balancing selection, split by journal.

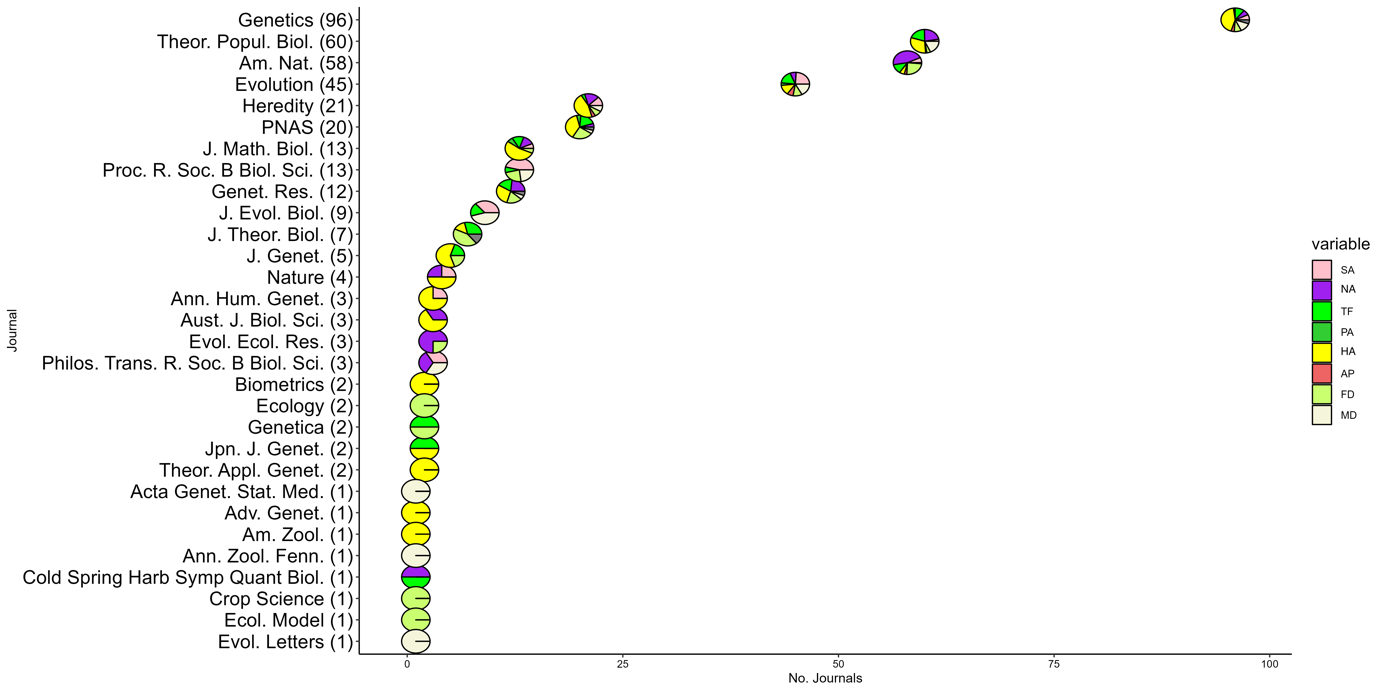

Timeline of publication of theoretical papers on balancing selection, split by journal and by scenario of balancing selection (SA = sexually antagonistic selection, NA = niche antagonism; TF = temporally fluctuating selection, PA = ploidally antagonistic selection, HA = heterozygote advantage, AP = antagonistic pleiotropy, FD = negative frequency-dependent selection, MD = meiotic drive).

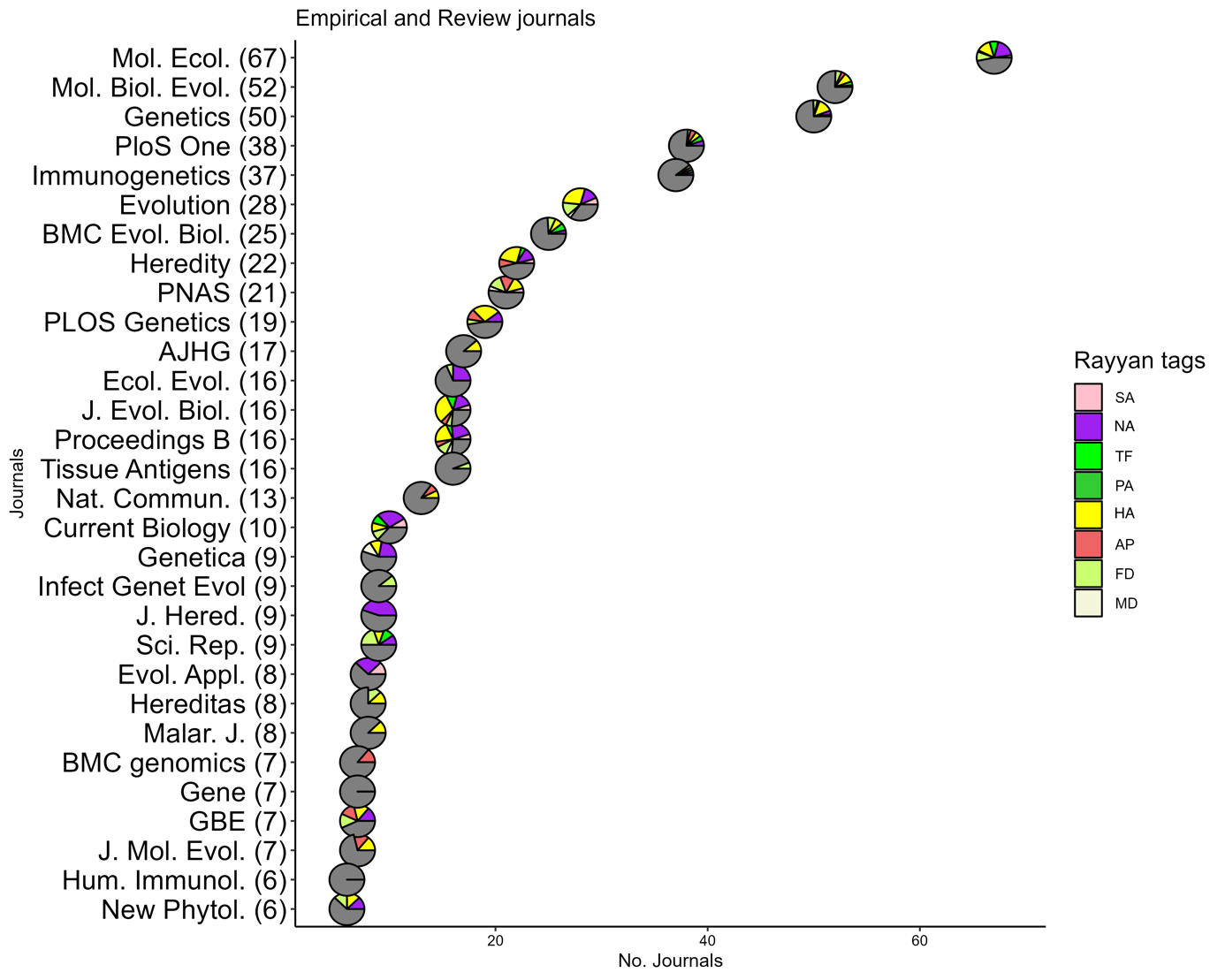

Timeline of publication of empirical papers on balancing selection, split by journal and by scenario of balancing selection (SA = sexually antagonistic selection, NA = niche antagonism; TF = temporally fluctuating selection, PA = ploidally antagonistic selection, HA = heterozygote advantage, AP = antagonistic pleiotropy, FD = negative frequency-dependent selection, MD = meiotic drive).
